# RETRO-URETERAL DELIVERY OF AAV-KP VECTORS FOR RENAL GENE TRANSFER AND THE ASSESSMENT OF TRANSGENE REABSORPTION IN THE KIDNEY

**DOI:** 10.64898/2026.11.18.767076

**Authors:** Sergio Milagros, Nerea Santana-Gonzalez, Martin Bilbao-Arribas, Leire Neri, Beatriz Carte, África Vales, Miriam Mendez, Leszek Lisowski, Nerea Zabaleta, Gloria Gonzalez-Aseguinolaza, Rafael Aldabe

## Abstract

Monogenic disorders cause a substantial fraction of chronic kidney disease, yet no renal gene therapy exists, partly because adeno-associated virus (AAV) vectors transduce the kidney poorly after systemic delivery. Screening a barcoded library of 80 natural and chimeric AAV serotypes administered retrogradely into the mouse renal pelvis identified AAV-KP1, AAV-KP2 and AAV-KP3 as enriched in the injected kidney but not in the liver. Individual validation confirmed efficient transduction of the administered kidney, with vector genome copy numbers approaching hepatic levels, 50- to 500-fold above the contralateral kidney and far above AAV9. Using kidney-specific (KSPC) and ubiquitous (CMV) promoters, transduction was detected in collecting duct, distal convoluted tubule, proximal tubule, Bowman’s capsule, interstitial cells and podocytes, with cell-type coverage depending on the promoter. Unexpectedly, GFP and mCherry protein was abundant in proximal tubules of kidneys lacking transgene mRNA, including contralateral and intravenously dosed kidneys. RNA-FISH, vector genome quantification and serum mCherry indicate that this protein derives from circulating transgene product filtered and reabsorbed by the proximal tubule rather than from local transduction. Retrograde delivery of AAV-KP capsids therefore achieves broad renal epithelial transduction, and transgene mRNA must be assessed alongside protein if kidney-tropic capsids are to be ranked reliably.

## INTRODUCTION

Chronic kidney disease (CKD) presents a major global health challenge and imposes a substantial economic burden. Genomic technologies have transformed our understanding of its causes. While diabetes and hypertension are associated with the majority of adult renal disease, genetic factors have a substantial contribution. Up to 30% to 70% of children requiring kidney replacement therapy can have an underlying monogenic cause, as do approximately 10% to 20% of adult-onset CKD cases (1,2); taken together, thus monogenic disorders account for roughly 6.5% to 10% of all CKD cases. Recent studies further demonstrate that 17% to 25% of CKD patients with previously unknown etiology can now be successfully reclassified with a precise genetic diagnosis (3,4). However, this diagnostic progress has not been matched therapeutically. Most monogenic kidney diseases still lack a disease-modifying treatment; and management relies primarily on generic nephroprotection, with dialysis and transplantation as the endpoint for most patients. Thus, there is an urgent need to develop alternative, long-term effective treatments for these patients.

Adeno-associated virus (AAV) vectors have become the leading platform for in vivo gene therapy, as reflected by the expanding array of approved AAV-based treatments and the large number of ongoing clinical trials (5,6). Nine products addressing monogenic hematologic have received approval to date (hemophilia A and B — Roctavian (7), Hemgenix (8), Beqvez (9)), neuromuscular (spinal muscular atrophy — Zolgensma (10); Duchenne muscular dystrophy — Elevidys (11)), neurometabolic (aromatic L-amino acid decarboxylase deficiency — Upstaza/Kebilidi (12)), retinal (Leber’s congenital amaurosis type 2 — Luxturna (13)), and hearing (Otoferlin-related deafness – Otarmeni (14)) disorders. In addition, numerous AAV gene therapy trials remain active across a range of organs and indications.

The route of administration largely reflects the accessibility of the target tissue. Systemic intravenous infusion is used for hepatic and neuromuscular disorders, as well as for hematologic disorders that exploit the liver as a protein factory. By contrast, ophthalmologic and otologic conditions require local surgical delivery bypassing the physical barriers that otherwise limit efficient vector access (6). For CNS-targeted therapies, both systemic and local administration strategies are used. Zolgensma is systemically deliver to reach motor neurons, albeit at high vector doses, while most brain directed therapies require intraparenchymal or intrathecal delivery which has motivated AAV capsid engineering capable of crossing the blood-brain barrier following intravenous administration (15).

Compared with other major organs, the use of genetic therapies to treat renal disease remains markedly underdeveloped. This gap is particularly significant given the substantial incidence of genetic causes underlying renal disease, many of which could in principle be addressed through gene transfer approaches.

Although no gene or cell therapies targeting the kidney have been approved yet, several experimental molecular treatments had been or are under evaluation. Teprasiran, a p53-directed siRNA relying on unconjugated proximal tubular uptake developed for the prevention of acute kidney injury (16), reached phase 3 but was discontinued after missing its primary endpoint (NCT03510897). The results obtained with RGLS8429 (Regulus Therapeutics), an anti-miR-17 oligonucleotide targeting the nephron epithelium, are more encouraging, having completed a phase 1b multiple ascending dose trial in Autosomal Dominant

Polycystic Kidney Disease patients with positive findings across dose cohorts (NCT05521191) (17). These therapies, however, require repeated lifelong administration, and nephrotoxicity from oligonucleotide accumulation in the proximal tubule is a recognized class effect (18,19). AAV gene therapy offers an alternative: a single administration with the potential for durable correction of monogenic renal disease.

The kidney nevertheless presents unique hurdles for targeted AAV-mediated gene delivery. It is a highly structured organ containing a diverse range of cell types along the renal epithelium. Each cell type executes distinct physiological and metabolic functions, and their dysfunction due to genetic causes gives rise to various renal diseases. Its intricate anatomical structure and glomerular filtration barrier restrict vector access, and systemic injections yields minimal transduction of most of nephron cell types even at high doses. Recently, cross-species directed evolution produced AAV.k13 and AAV.k20, which transduce murine proximal tubules after IV dosing more efficiently that the parental AAV9 serotype (20). However, local renal administration significantly improves epithelium transduction, this approach bypasses the glomerular filtration barrier and allows the vectors to reach the renal epithelium directly, reducing vector dissemination (20,21). Several AAV serotypes have been shown to transduce renal cells in mice, swine and non-human primates being, in all the cases, the proximal tubular cells the cell type more efficiently transduced (20–22). Notably, other groups have reported glomerular transduction with selected AAV capsids following both systemic and local delivery (23–25). The local route has advanced furthest, reaching the clinic in a phase 1/2 trial for primary IgA nephropathy (NCT07182227).

Thus, the development of AAV-based therapies for genetic kidney disease therefore requires a systematic characterization of renal transduction, establishing which nephron cell types can be efficiently targeted and which diseases are consequently amenable to treatment. With this aim, we screened an AAV library composed of 80 natural and chimeric serotypes, administered retrogradely, to identify vectors with higher renal tropism. We validated the renal tropism of AAV-KP1, AAV-KP2, and AAV-KP3, demonstrating transduction of different renal epithelial cell types in addition to interstitial and glomerular cells. Importantly, we show that transgene mRNA and protein must be assessed together, since protein detection alone does not establish that the cell in which it is visualized is the cell that produced it. We thereby identify a reabsorption-driven artifact and define the controls required to measure renal transduction correctly. This artifact plausibly accounts for part of the heterogeneity in reported proximal tubule efficiencies after AAV vector administration, and has important consequences for how candidate kidney-tropic capsids are ranked. Moreover, this transgene protein reabsorption might have implications for the evaluation of the therapeutic activity of gene therapy vectors for renal disorders.

## MATERIALS AND METHODS

### AAV vector production and purification

AAV vectors were produced in HEK293T cells. For each production, cells were expanded across sixty 150-mm plates and transfected at approximately 70% confluency. Transfection was performed using polyethylenimine (PEI) at a 1:1 DNA:PEI ratio, with 40 μg pDF6 helper plasmid, 20 μg capsid plasmid and 20 μg ITR-flanked expression plasmid per plate (26).

Cells and supernatant were harvested separately 72 h after transfection. Cells were resuspended in 1× TMS lysis buffer (0.5 M Tris, 1.5 M NaCl, 20 mM MgCl₂, 1% Triton X-100) and subjected to three freeze–thaw cycles to release viral particles. Vector in the supernatant was precipitated with 40% PEG in 2.5 M NaCl (48–72 h, 4 °C) and the precipitate was resuspended in 1× TMS lysis buffer. Pooled lysates were treated with DNase I and RNase A (Roche, Basel, Switzerland; cat. nos. 10104159001 and 10109169001; 0.1 mg per 10 μL) for 60 min at 37 °C to remove non-encapsidated nucleic acids.

Lysates were purified on a discontinuous 15–57% iodixanol gradient (OptiPrep Density Gradient Medium, cat. no. D1556; Sigma-Aldrich, St Louis, MO, USA). Following ultracentrifugation, AAV particles were recovered from the 40–57% interphase and buffer-exchanged into PBS containing 5% sucrose using Amicon Ultra-15 centrifugal filters (cat. no. UFC9100; Merck, Darmstadt, Germany) at 4000 × *g*. Vectors were further purified by affinity chromatography on an ÄKTA start system (GE Healthcare Life Sciences, Uppsala, Sweden) using POROS GoPure AAVX pre-packed 1-mL columns (cat. no. A36652; Thermo Fisher Scientific, Waltham, MA, USA). Columns were equilibrated in PBS (pH 7.4), loaded with sample, washed sequentially with PBS and PBS containing 1 M NaCl, and eluted with 50 mM citric acid (pH 3.0). Eluates were neutralised immediately with 1 M Tris (pH 9.0), concentrated using Amicon Ultra-15 filters, sterile-filtered through a 0.45-μm membrane and stored at −80 °C.

### Animals manipulation

Male and female C57BL/6J mice were purchased from Charles River Laboratories (Saint-Germain-Nuelles, France) and housed under specific pathogen-free conditions on a 12-h light/dark cycle, with *ad libitum* access to food and water. All experimental procedures were approved by the Ethics Committee for Animal Testing of the University of Navarra (protocol 069-20 and 117-25) and conducted in accordance with European Directive 2010/63/EU and Spanish Royal Decree 53/2013. Reporting follows the ARRIVE guidelines.

Six-week-old mice received a single administration of the corresponding AAV vector by either the intravenous or the retro-ureteral route. Animals were euthanised two weeks after vector administration. Before perfusion, blood was collected by submandibular vein puncture; serum was separated by centrifugation (4 °C) and stored at −80 °C for subsequent ELISA. Mice were then anaesthetised with a ketamine (Richter Pharma)/xylazine (Rompun) mixture and transcardially perfused with 10 mL PBS injected into the left ventricle, with a right atrial incision as the outflow. Left and right kidneys and liver were collected and processed for RNA extraction, genomic DNA (gDNA) extraction and immunohistochemistry (IHC).

#### Intravenous administration

Mice were anaesthetised with isoflurane (IsoVet, B. Braun, Melsungen, Germany) and given a single retro-orbital injection of the corresponding AAV vector at 1 × 10¹⁰ vg per gram body weight, diluted in PBS containing 5% sucrose to a final volume of 100 µL.

#### Retro-ureteral administration

Mice were anaesthetised by isoflurane inhalation and placed supine on a rodent surgical table; the abdomen was shaved and disinfected with povidone-iodine. A midline laparotomy was performed slightly left of the midline, and the intestines were retracted within the abdominal cavity to expose the left kidney and ureter. The renal artery, renal vein and ureter were temporarily occluded with micro-serrefine clamps (Fine Science Tools, Heidelberg, Germany; #18055-04) to interrupt blood flow and urine outflow before vector delivery. A 30G needle connected to a 1 mL syringe mounted on a syringe pump (Legato 130 Nanoliter, KD Scientific, Holliston, MA, USA; #788130) was inserted into the renal pelvis, and 100 µL of AAV vector solution was infused at a controlled flow rate of X µL/min. Clamps were maintained for a total of 15 min to enhance transduction, and the surgical field was kept moist with sterile saline throughout. After clamp release, restoration of renal blood flow was confirmed visually and the organs were repositioned. The abdominal wall and skin were closed with 6-0 sutures (Ethicon, #ETV489), and buprenorphine (Bupaq,) was given for post-operative analgesia. Animals recovered on a heating pad until fully ambulatory.

### Vector genome quantification

Genomic DNA (gDNA) was extracted from tissue with the Maxwell RSC Tissue DNA Kit (Promega, #AS1610) using a modified protocol. DNA concentration and purity were assessed spectrophotometrically.

Vector genome copy number was determined by quantitative PCR using iQ SYBR Green Supermix (Bio-Rad, Hercules, CA, USA). Reactions were run in triplicate, and melt-curve analysis was performed to confirm amplification specificity.

Transgene (*GFP*) and promoter (*CMV*) amplicons were quantified and normalised to the single-copy endogenous gene *Rb36b4* by the ΔCt method, with relative vector genome content calculated as 2^(Ct_reference − Ct_target)^. Primer sequences are:

- *GFP:* Forward (5′→3′) GACGACGGCAACTACAAGAC; Reverse (5′→3′) TCCTTGAAGTCGATGCCCTT.
- *CMV:* Forward (5′→3′) TTCAACATGGTGATGCGGTT; Reverse (5′→3′) TACACGCCTTACCCCGCATT.
- *Rb36b4:* Forward (5′→3′) AACAACTCCGCCCTCTCCTT; Reverse (5′→3′) GAAGGCCTTGACCTTTCACG.

### Gene expression quantification

Total RNA was extracted from cells and tissues using the Maxwell RSC simplyRNA Tissue Kit (Promega, Madison, WI, USA; #AS1340) on a Maxwell RSC 48 instrument, following a modified version of the manufacturer’s protocol. RNA concentration and purity were assessed by spectrophotometry. Genomic DNA was removed by treating 1 µg of RNA with DNase I (1 U) in the presence of RNaseOUT (16 U), followed by heat inactivation at 65 °C for 10 min and RNA denaturation at 90 °C for 1 min. First-strand cDNA was synthesised with M-MLV reverse transcriptase (120 U) and random hexamers (200 ng; Roche, Basel, Switzerland) for 1 h at 37 °C, followed by enzyme inactivation at 95 °C for 1 min. Quantitative PCR was performed with iQ SYBR Green Supermix (Bio-Rad, Hercules, CA, USA), using primers specific for *GFP*, *mCherry* and *Rb36b4*. *GFP* and *mCherry* expression were normalised to *Rb36b4* and calculated using the 2^−ΔΔCt method. All reactions were run in triplicates. Primer sequences are:

- *GFP:* Forward (5′→3′) ATGGTGAGCAAGGGCGAGGA; Reverse (5′→3′) TTCCCGGTGGTGCAGATGAA.
- *mCherry:* Forward (5′→3′) GAGATCAAGCAGAGGCTGAA; Reverse (5′→3′) ATGGTGTAGTCCTCGTTGTG.
- *Rb36b4:* Forward (5′→3′) AACAACTCCGCCCTCTCCTT; Reverse (5′→3′) GAAGGCCTTGACCTTTCACG.

### Histology, immunohistochemistry and immunofluorescence

Tissue samples were fixed in 4% paraformaldehyde (Panreac) for 48 h at room temperature, transferred to 70% ethanol, paraffin-embedded and sectioned at 3 µm by the Histology Core Facility at CIMA (University of Navarra, Pamplona, Spain). Sections were stained with haematoxylin and eosin for histological assessment, or processed for immunohistochemical (IHC) and immunofluorescence (IF) detection. Sections were blocked and incubated with the primary antibodies: anti-THP (Santa Cruz Biotechnology, sc-271022), anti-AQP2 (Santa Cruz Biotechnology, sc-515770), anti-eGFP (Novus Biologicals, NB100-1770) and anti-mCherry (Invitrogen, M11217) according to the manufacturer’s instructions for each antibody and using the appropriate detection systems. Nuclei were counterstained with haematoxylin (IHC) or DAPI (IF). Negative controls were processed in parallel with omission of the primary antibody. IHC images were acquired on a Leica Aperio CS2 scanner using Aperio ImageScope software and the IF images were acquired on a Zeiss LSM 800 laser scanning confocal microscope using ZEN software.

### RNAscope

RNAscope fluorescent *in situ* hybridisation was performed on 5 µm paraffin-embedded sections using the RNAscope Multiplex Fluorescent Reagent Kit v2 (Bio-Techne/ACD, Newark, CA, USA; #323100) following the manufacturer’s protocol. Independent sections from each sample were used to detect *mCherry* and *eGFP* transcripts. Sections were deparaffinised, subjected to target retrieval and protease treatment, and hybridised separately with custom probes against *mCherry* (#431201) or *eGFP* (#400281). Probe signal was amplified using the HRP-based amplification reagents supplied with the kit and developed with TSA Vivid Opal 570 (#323272). Nuclei were counterstained with DAPI (#320858).

Positive and negative control probes were processed in parallel on adjacent sections. Slides were scanned on a Vectra Polaris Imaging System. Acquisition parameters were held constant within each probe set (*mCherry*: DAPI 6.64 ms, Opal 570 83 ms; *eGFP*: DAPI 6 ms, Opal 570 250 ms) but differed between probe sets; signal intensities are therefore not directly comparable between the two.

### Quantification of serum mCherry by ELISA

Serum mCherry was quantified by sandwich ELISA (mCherry ELISA Kit, Abcam, #ab325452) according to the manufacturer’s instructions. Samples were diluted in the supplied Sample Diluent and assayed in duplicate. All samples were analysed at 1:250, together with a second dilution selected according to the expected mCherry concentration: 1:100 for samples with lower expected levels and 1:500 for those with higher expected levels, so that each concentration was determined from two independent dilutions falling within the assay range.

An eight-point standard curve was freshly prepared for each assay by two-fold serial dilution (400 to 6.25 pg/mL) and included a blank. Standards and samples (50 µL, in duplicate) were combined with 50 µL of capture/detection antibody cocktail per well (100 µL final volume) and incubated for 1 h at room temperature with shaking at 400 rpm. Plates were washed three times with Wash Buffer, developed with TMB substrate for 10 min in the dark with shaking, and the reaction was stopped and absorbance was read at 450 nm. Blank-corrected absorbance values were fitted to the standard curve by four-parameter logistic (4PL) regression model. Serum mCherry concentrations were derived from the curve and corrected for the corresponding dilution factor.

### AAV library sequencing and barcode quantification

Total RNA was extracted from tissue samples using the AllPrep DNA/RNA Mini Kit (QIAGEN; #80204). RNA was treated with the TURBO DNA-*free* Kit (Invitrogen; #AM1907) to remove residual gDNA. First-strand cDNA was synthesised with the SuperScript IV First-Strand Synthesis System (Invitrogen; #18091050) using a gene-specific reverse primer.

Barcode regions were amplified from cDNA and gDNA using Q5 Hot Start High-Fidelity DNA Polymerase (New England Biolabs, #M0493) and dNTP Solution Mix (NEB; #N0447), with sample-specific barcoded forward primers and a universal reverse primer:

- BC_0039_GFP_BC_01 5′: GTTCAgctggagttcgtgaccgccg-3’
- BC_0040_GFP_BC_02 5′: GTCATgctggagttcgtgaccgccg-3′
- BC_0041_GFP_BC_03 5′: CTGTAgctggagttcgtgaccgccg-3′
- Universal reverse primer: 5′-CAACATAGTTAAGAATACCAGTCAATCTTTCAC-3′

PCR products were resolved on 2% agarose gels (Sigma-Aldrich, #A9539) in 1× TAE buffer (Life Technologies, #24710-030) alongside a 1 kb Plus DNA Ladder (NEB; #N3200). Amplicons of the expected size were purified with the Zymoclean Gel DNA Recovery Kit (Zymo Research, #D4007), and pooled in equimolar amounts before library preparation.

Library preparation and paired-end sequencing (2 × 150 bp) were performed by the Genomics Core Facility at CIMA on an Illumina platform (Illumina, San Diego, CA, USA). Reads were randomly subsampled from an initial depth of approximately 100 million to 10 million read pairs per sample to reduce computational load. Raw reads were quality-filtered and adapter-trimmed with bbduk.sh (BBTools v39.01; https://sourceforge.net/projects/bbmap), merged into single reads with bbmerge.sh (27) and filtered by expected amplicon length with bbduk.sh. Demultiplexing and quantification of tissue- and sample-specific counts across the 81-variant AAV library were performed with a custom Python script that identifies barcode combinations at defined positions (https://doi.org/10.24433/CO.7176285.v2) (28). Raw counts were normalised to total counts per tissue and expressed as a percentage.

## RESULTS

### Identification and characterization of AAV serotypes with renal tropism following retro-ureteral administration

Few AAV vectors have been described for renal targeting. To identify serotypes with renal tropism, we administered a barcoded library of 80 natural and chimeric AAV serotypes expressing GFP into the left renal pelvis of mice, after clamping the renal artery, renal vein, and ureter (Figure 1A). Two weeks later, we harvested liver, left kidney, and right kidney samples and analyzed them for serotype-specific barcodes to identify those transducing the renal epithelium.

**Figure 1.**
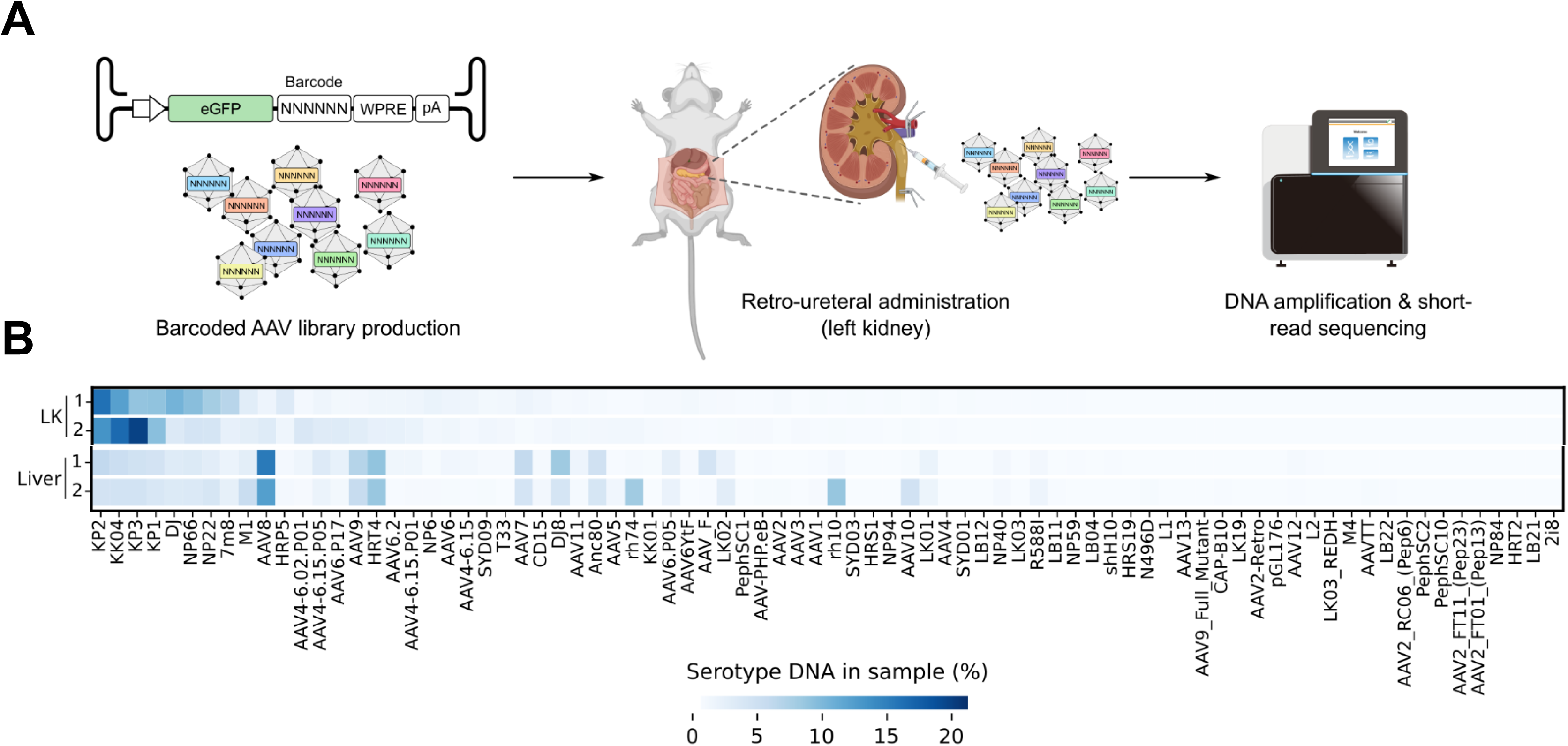
Identification of AAV serotypes with renal tropism. **(A)** Eight-week-old mice received a retrograde injection into the left renal pelvis of 5 × 10¹⁰ vg/g of a barcoded AAV library comprising 80 capsid variants, each carrying a unique barcode positioned between the GFP open reading frame and the WPRE–polyA sequence. Two weeks after administration, mice were euthanised and barcode representation in the injected kidney and in the liver was determined by cDNA amplification and short-read sequencing. **(B)** Relative abundance of each AAV capsid in the injected left kidney (LK) and in the liver, expressed as a percentage of total barcode counts and shown on a blue colour scale.

Sequencing revealed a clear enrichment of serotypes AAV-KP1, AAV-KP2, AAV-KP3, and AAV.KK04 in the injected kidney (Figure 1B). Interestingly, the serotypes most represented in the liver included AAV8, AAV9, and AAV7, which have well-described hepatic tropism in mice (29). In contrast, none of the renal tropic serotypes was among the most represented serotypes in the liver (Figure 1B).

Next, we produced the four AAVs with the best renal profile — AAV-KP1, AAV-KP2, AAV-KP3 and AAV.KK04 — each carrying the same expression cassette that contains the GFP coding sequence driven by the kidney-specific cadherin (KSPC) promoter (30). Each vector was individually administered into the renal pelvis of nine to ten mice; however, due to the technical complexity of the surgical procedure, a subset of animals could not be successfully dosed and was excluded from the analysis. Two weeks after injection mice were sacrificed to evaluate transduction efficacy.

Vector genome copy numbers in the administered (left) kidney were highest with AAV-KP1 and AAV-KP3, intermediate with AAV-KP2, and lowest with AAV.KK04 (Figure 2A and Supplementary Figure 1A). In contrast AAV-KP2 was the most efficient transducing the liver and the non-injected right kidney, AAV-KP1 and AAV-KP3 result in comparable and substantially lower levels, and AAV-KK04 again gave the lowest (Figure 2A and Supplementary Figure 1A). For all four serotypes, the non-administered contralateral kidney contained 50-to 500-fold fewer genome copies than the administered kidney.

**Figure 2.**
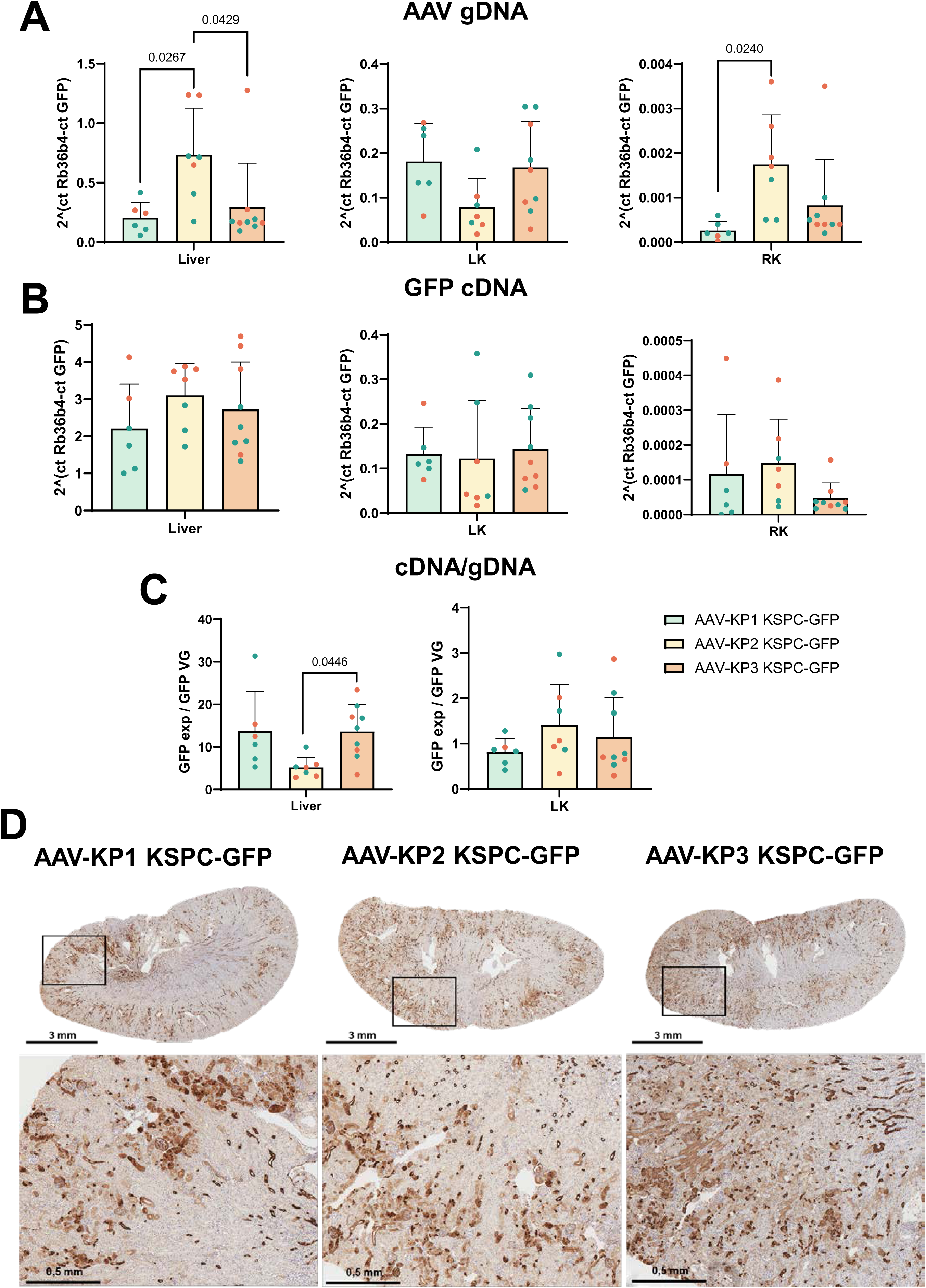
Renal and hepatic transduction efficiency of AAV-KP1, AAV-KP2 and AAV-KP3 after retro-ureteral administration. Six-week-old mice received 1 × 10¹⁰ vg/g of AAV-KP1, AAV-KP2 or AAV-KP3 vectors carrying GFP under the KSPC promoter, injected into the left renal pelvis. Mice were euthanised two weeks later to quantify vector genomes and GFP transgene expression. **(A)** Vector genome (vg) copies in the liver and both kidneys two weeks after administration, measured by quantitative PCR. **(B)** GFP transcript levels in the liver and both kidneys two weeks after administration, measured by RT-qPCR. **(C)** GFP mRNA normalised to vector genome copy number. **(D)** GFP-positive cells in renal sections detected by immunohistochemistry. The boxed region in each image is shown at higher magnification below. Scale bars are indicated in each image. Green symbols denote male and orange symbols female animals. Statistical analysis of the groups in each case was performed using Tukey’s multiple comparisons test. Numerical values indicates the p values comparing every group. Mean + SD was represented

Overall, transgene expression levels broadly correlated with the presence of vector genomes: in the administered kidney, AAV-KP1 and AAV-KP3 showed the highest transgene expression, AAV-KP2 did not differ significantly from these, although the majority of the animals showed lower expression levels (Figure 2B). In the liver, the highest expression levels were observed in the AAV-KP2 group. Contrary to expectation, in the context of the AAV genome the KSPC promoter showed higher transcriptional activity in liver than in kidney, measured as GFP mRNA per vector genome (Figure 2C), revealing a lack of specificity not predicted by earlier characterisations of this promoter.

Notably, when data from all AAV-injected groups were pooled, we observed that renal transgene expression was significantly higher in male than in female mice while hepatic expression was higher in female mice (Supplementary figure 1B).

Then GFP protein expression was determine by immunohistochemistry (IHC) analysis in renal sections of animals dosed with AAV-KP1, AAV-KP2 and AAV-KP3, observing a similar expression pattern with all three vectors (animals dosed with AAV.KK04 were not included in this analysis because of the low transduction efficiency of this serotype) (Figure 2D). In the administered (left) kidney, transduction was observed across different renal cell types in both the medulla and cortex, including collecting duct, distal convoluted tubule, proximal tubule and Bowman’s capsule (Figure 2D). In the right kidney, only a few scattered GFP-positive cells were observed throughout the renal parenchyma across the three experimental groups. Interestingly, three females administered with AAV-KP2 (RK2, RK3, RK4) and two females administered with AAV-KP3 (RK1, RK4) showed numerous proximal tubular cells with apical GFP-positive staining, a pattern also observed in their corresponding left kidneys (Supplementary Figure 2). Notably, although these right kidneys showed clear GFP immunostaining, GFP transcript levels were low and comparable to those in the right kidneys of the other mice across the three AAV groups, revealing a discordance between transcript and protein detection.

### Comparative renal transduction of AAV-KP1, AAV-KP3 and AAV9 carrying GFP under the transcriptional control of a ubiquitous expression promoter

To confirm and further characterize the observed renal tropism of the selected capsids, we administered into the left renal pelvis AAV-KP1 and AAV-KP3 expressing GFP under the control of the ubiquitous constitutive cytomegalovirus (CMV) promoter. This was also done to determine whether additional renal cell types were transduced by these AAVs but not detected due to lack of activity of the KSPC promoter in those cell types. In parallel, we administered an AAV9 carrying the same expression cassette, as it has been described as a renal-tropic vector by other authors (31,32).

As described above, both KP serotypes achieved similar transduction levels in the injected kidney, whereas animals receiving AAV9 showed 60-to 95-fold fewer vector genomes (Figure 3A). The contralateral non-injected kidney showed comparable AAV genome levels across all three serotypes and evaluation of AAV genome presence showed similar hepatic transduction among all three AAV serotypes. Accordingly, transgene expression in each organ reflected the amount of vector genomes present, with higher renal transgene expression for AAV-KP1 and AAV-KP3 than for AAV9 (Figure 3B, Supplementary Figure 3). Interestingly, transgene expression in the liver was similar to renal expression in AAV-KP1 and AAV-KP3 treated mice (Figure 3B).

**Figure 3.**
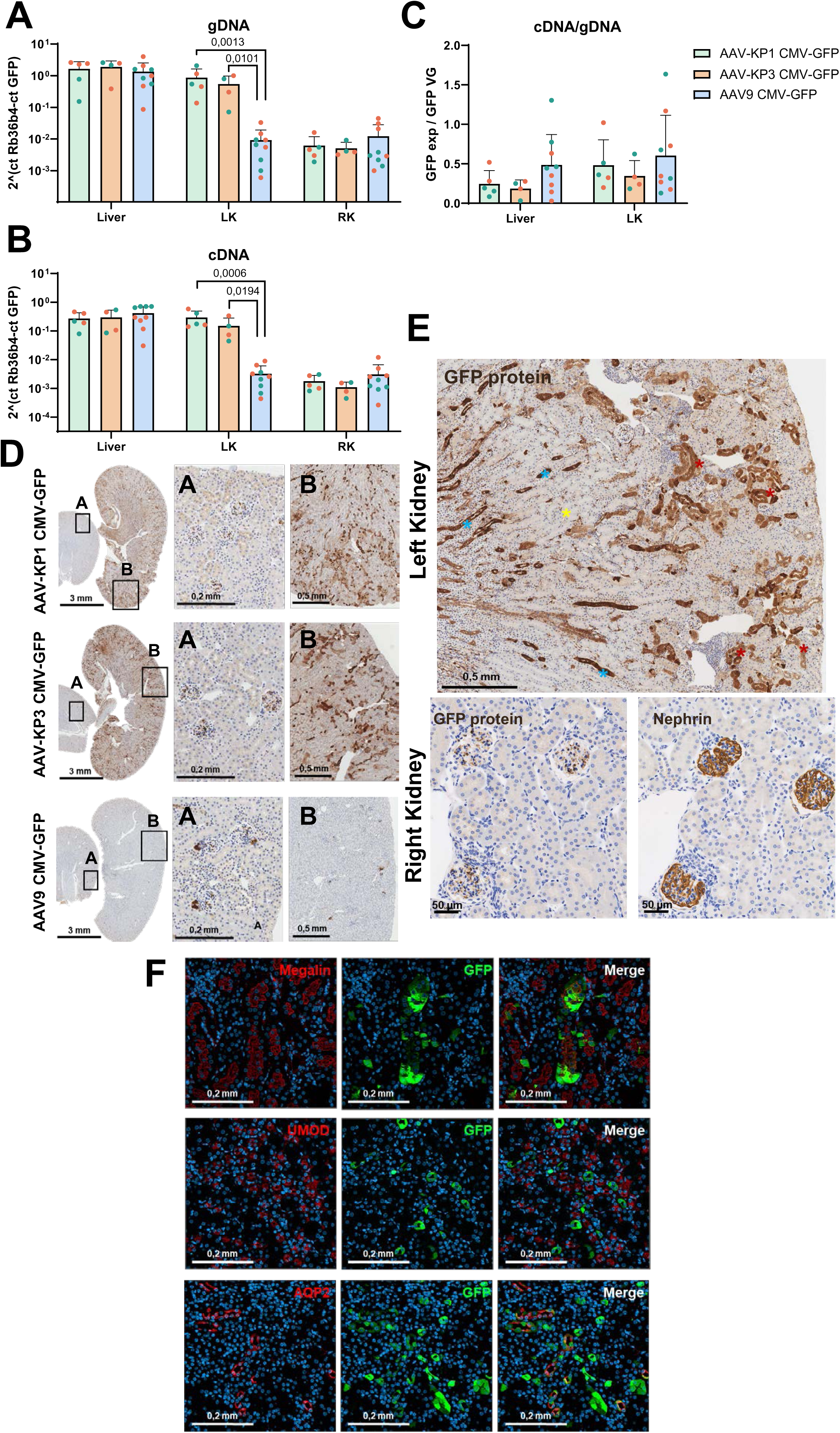
Renal and hepatic transduction efficiency of AAV-KP1, AAV-KP3 and AAV9 CMV-GFP vectors after retro-ureteral administration. Six-week-old mice received 1 × 10¹⁰ vg/g of AAV-KP1, AAV-KP3 or AAV9 vectors carrying GFP under the CMV promoter, injected into the left renal pelvis. Mice were euthanised two weeks later to quantify vector genomes and GFP transgene expression. **(A)** Vector genome (vg) copies in the liver and both kidneys, measured by quantitative PCR. **(B)** GFP transcript levels in the liver and both kidneys, measured by RT-qPCR. **(C)** GFP mRNA normalised to vector genome copy number. **(D)** GFP-positive cells detected by immunohistochemistry in sections of the right (upper panels) and left (lower panels) kidneys. Boxed regions are shown at higher magnification. Scale bars are indicated in each image. **(E)** Renal cell types transduced in the left kidney after administration of AAV-KP1 CMV-GFP: distal convoluted tubule (**\***), collecting duct (**\***) and interstitial cells (**\***). Podocytes (nephrin) and GFP-positive cells are shown in serial sections of the right kidney. **(F)** Confocal detection of GFP-positive cells (green) together with markers of renal epithelial subpopulations (red): megalin (proximal tubule), uromodulin (thick ascending limb of the loop of Henle) and aquaporin-2 (collecting duct). Green symbols denote male and orange symbols female animals. Data are mean ± SD. Groups were compared using Tukey’s multiple comparisons test; numerical values indicate P values for each pairwise comparison.

Consistent with the renal cellular tropism described above, AAV-KP1 and AAV-KP3, transduced several renal epithelial cell types, including proximal tubule, distal convoluted tubule and collecting duct (Figures 3C, 3D and 3E); however, with the CMV promoter GFP was additionally detected in interstitial cells and podocytes, but not Bowman’s capsule cells that were detected with the KSPC promoter. No GFP-expressing cells were detected in the thick ascending limb of the loop of Henle, nonetheless, taking into consideration the differences observed depending on the promoter we cannot rule out that the absence of transgene expression in both studies reflects promoter activity rather than a lack of transduction. Thus, multiple promoters should be used to identify the full range of transduced renal cell types.

Interestingly, AAV9-treated mice showed a very similar renal transduction pattern in both kidneys, with positive cells in juxtaglomerular apparatus and glomeruli, likely podocytes as it has been described previously (23).

Notably, although KSPC promoter is described as kidney-specific, the 1.5 kb KSPC fragment used here drove strong hepatic expression, approximately 10-fold higher than that achieved with the CMV promoter (Supplementary Figure 4).

### Influence of the administration route on renal transduction

We next compare the biodistribution of AAV-KP1 CMV-GFP administered retrograderaly in the renal pelvis or intravenously. Retro-ureteral administration of AAV-KP1 efficiently transduced the left kidney, at levels comparable to the liver (Figure 4). In contrast, systemic administration resulted in hepatic transduction together with substantially lower and similar transduction levels in both kidneys, comparable to those observed in the right (non-administered) kidney following retro-ureteral delivery to the left kidney (Supplementary Figure 5). As expected, systemic administration produced substantially higher hepatic transduction than the retro-ureteral route (Figure 4C).

**Figure 4.**
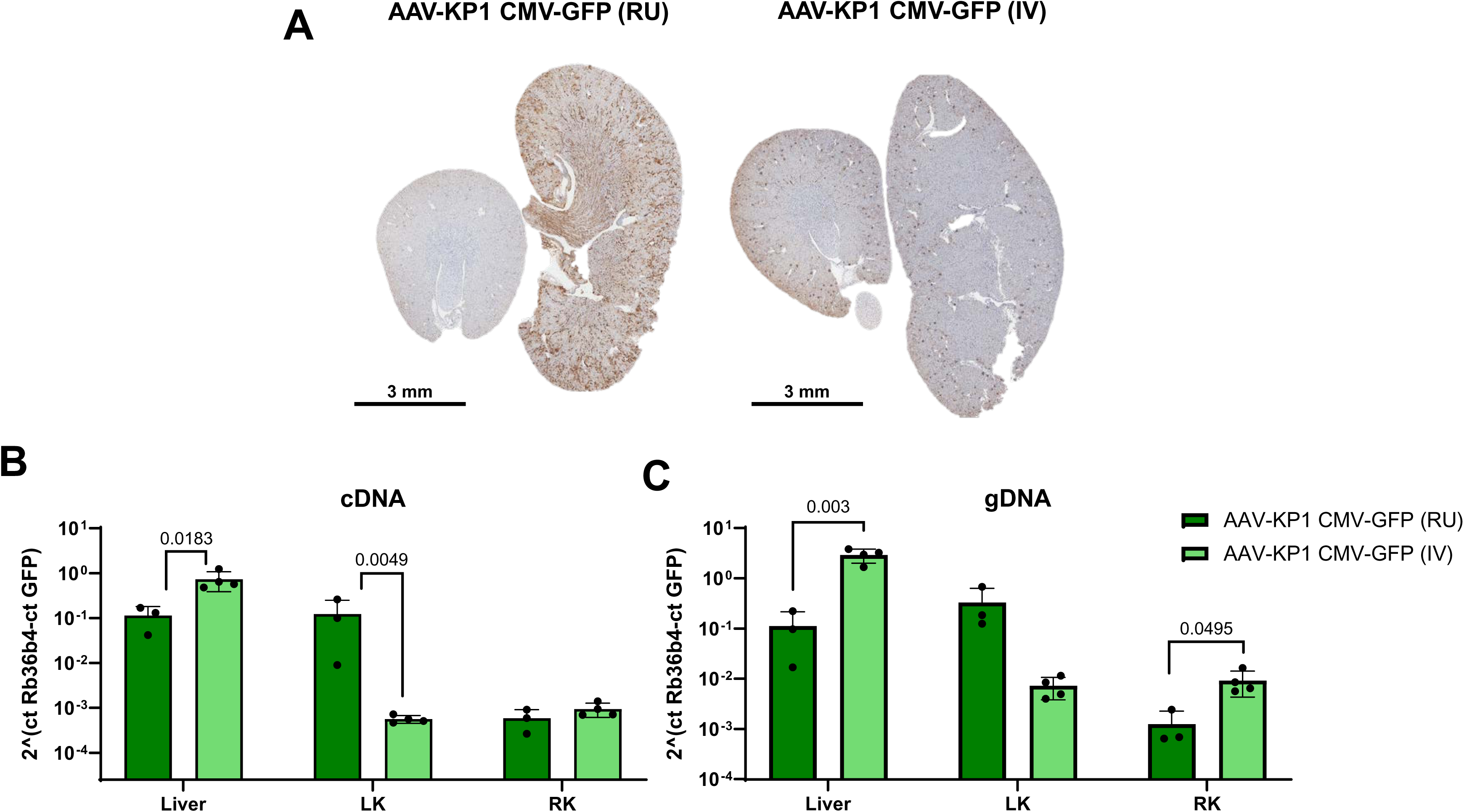
Renal transduction efficiency of the AAV-KP1 CMV-GFP vector after systemic or retro-ureteral administration. Six-week-old mice received 1 × 10¹⁰ vg/g of the AAV-KP1 vector carrying GFP under the CMV promoter, delivered either into the left renal pelvis (RU) or intravenously (IV). Mice were euthanised two weeks later to quantify vector genomes and GFP transgene expression. **(A)** GFP-positive cells detected by immunohistochemistry in sections of the right (upper kidney) and left (lower kidney) kidneys. Boxed regions are shown at higher magnification. Scale bars are indicated in each image. **(B)** GFP transcript levels in the liver and both kidneys, measured by RT-qPCR. **(C)** Vector genome (vg) copies in the liver and both kidneys, measured by quantitative PCR. Data are mean ± SD. Groups were compared using Tukey’s multiple comparisons test; numerical values indicate P values for each pairwise comparison.

Similar results were obtained when the reporter was changed to mCherry (Figure 5, Supplementary Figure 6). Mice received AAV-KP1 CAG-mCherry either renal retrograde route or intravenously, together with an additional group receiving AAV9 CAG-mCherry intravenously.

**Figure 5.**
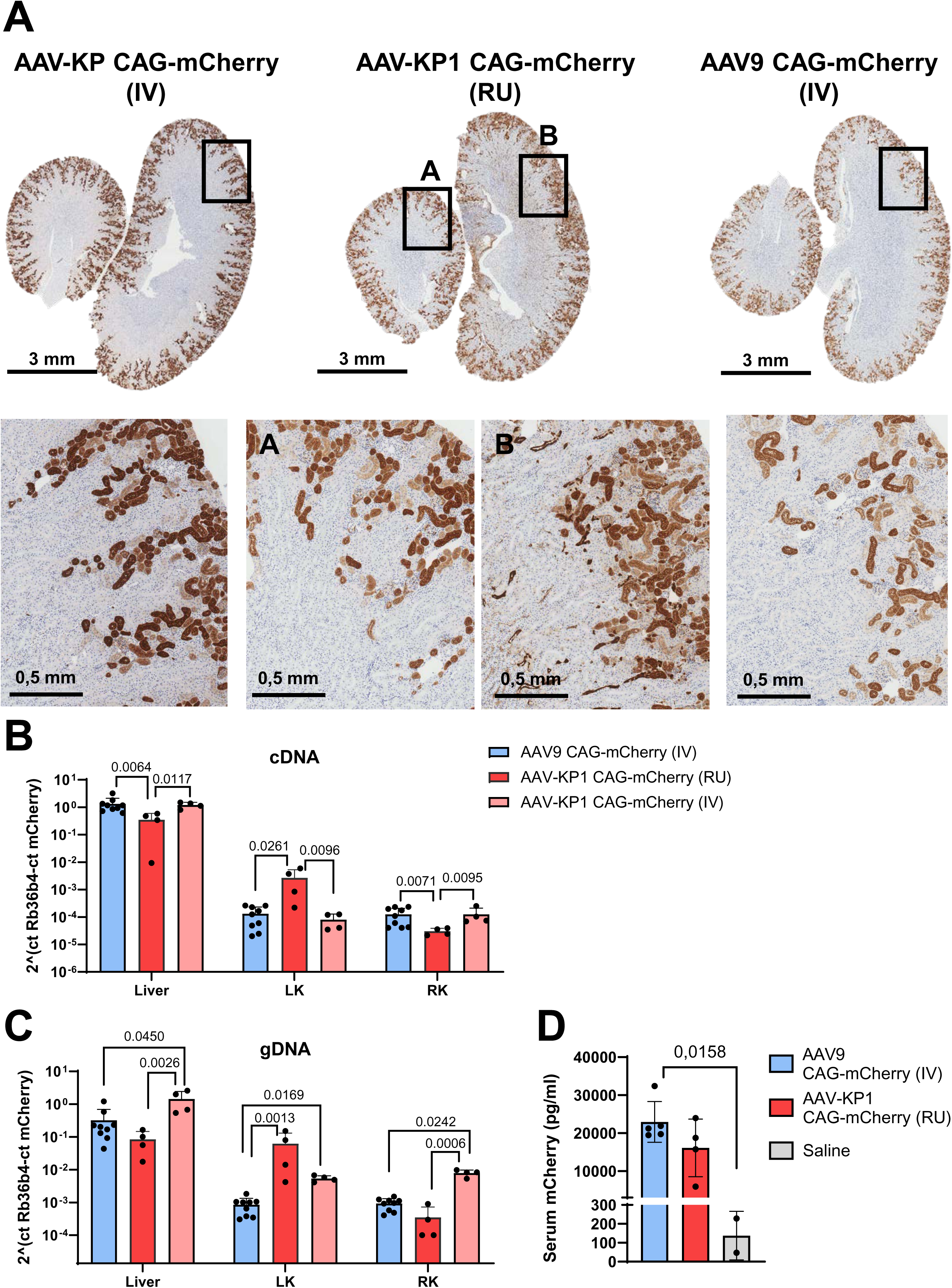
Renal transduction efficiency of the AAV-KP1 CAG-mCherry vector after systemic or retro-ureteral administration and AAV9 CAG-mCherry vector after systemic administration. Six-week-old mice received 1 × 10¹⁰ vg/g of the AAV-KP1 or AAV9 vector carrying mCherry under the CAG promoter, delivered either into the left renal pelvis (RU) or intravenously (IV). Mice were euthanised two weeks later to quantify vector genomes and mCherry transgene expression. **(A)** mCherry-positive cells detected by immunohistochemistry in sections of the right (lower kidney) and left (upper kidney) kidneys. Boxed regions are shown at higher magnification. Scale bars are indicated in each image. **(B)** mCherry transcript levels in the liver and both kidneys, measured by RT-qPCR. **(C)** Vector genome (vg) copies in the liver and both kidneys, measured by quantitative PCR. **(D)** mCherry concentration in mice sera quantified by ELISA. Data are mean ± SD. Groups were compared using Tukey’s multiple comparisons test; numerical values indicate P values for each pairwise comparison.

Kidneys from all treated mice showed strong mCherry signal in the proximal tubule, with preferential apical staining of the renal epithelium, irrespectively of the route of administration. This pattern resembles that seen in a subset of animals in the preceding experiments (Supplementary Figure 7). Quantification of mCherry mRNA and vector genome copy number in renal and hepatic samples reproduced the observations made with the AAV-KP1 CMV-driven GFP vector, mCherry mRNA level in the retrograde administered left kidney approached hepatic levels, whereas mCherry expression was very low in the contralateral right kidney and in both kidneys of mice administered either vector intravenously. Genome copy number correlated with mCherry mRNA expression, consistent with the results obtained with the GFP-expressing vectors. Accordingly, the highest genome copy numbers were found in the liver of mice administered AAV-KP1 or AAV9 intravenously and, at similar levels, in the liver and left kidney of mice administered with AAV-KP1 retro-ureterally. In contrast, the right kidneys of all treated mice and the left kidneys of intravenously administered animals showed very low transduction. These results also revealed a promoter-dependent difference in renal activity. In retrogradely administered mice, genome copy numbers were comparable between the left kidney and the liver, yet mCherry expression driven by the CAG promoter was approximately 100-fold lower in the kidney than in the liver, whereas GFP expression driven by the CMV promoter was similar in both organs. The CAG promoter is therefore markedly less active in the treated kidney than in the liver, in contrast to CMV, which performs comparably across the two organs.

### Characterization of transgene presence in the renal tissue after AAV administration

Taking together, these data revealed an important inconsistency between transgene mRNA expression and the presence of GFP or mCherry protein by IHC in the right kidney of animals that received the vectors retro-ureterally in the left kidney (Figure 4 and 5, Supplementary Figure 5 and 6). GFP protein levels in the apical region of proximal tubule epithelial cells, were comparable in the two kidneys, whereas transgene mRNA in the right kidney was nearly 100-fold lower than in the left kidney.

To investigate this discrepancy between mRNA and protein, we performed RNA-FISH for GFP transcripts on renal and hepatic sections from mice administered with AAV9 CMV-GFP or AAV-KP1 CMV-GFP retro-ureterally, or AAV-KP1 CMV-GFP intravenously.

GFP mRNA was detected in the liver of all three groups and in the left kidney of AAV-KP1-CMV-GFP retro-ureterally injected mice. In contrast, in the kidneys of intravenously dosed animals and the right kidney of retro-ureterally injected mice, GFP transcript was detected only in the glomeruli (Figure 6A and Supplementary Figure 8). Similar results were obtained in the animals receiving mCherry expressing vectors (AAV9 CAG-mCherry or AAV-KP1 CAG-mCherry intravenously and AAV-KP1 CAG-mCherry retro-ureterally). In all three groups, mCherry transcript was detected in hepatic sections, as expected, and broad mCherry mRNA expression was detected in the left kidney sections from mice locally administered with AAV-KP1 CAG-mCherry (Figure 6B and Supplementary Figure 9). In all other renal sections, mCherry mRNA was detected only in the glomeruli.

**Figure 6.**
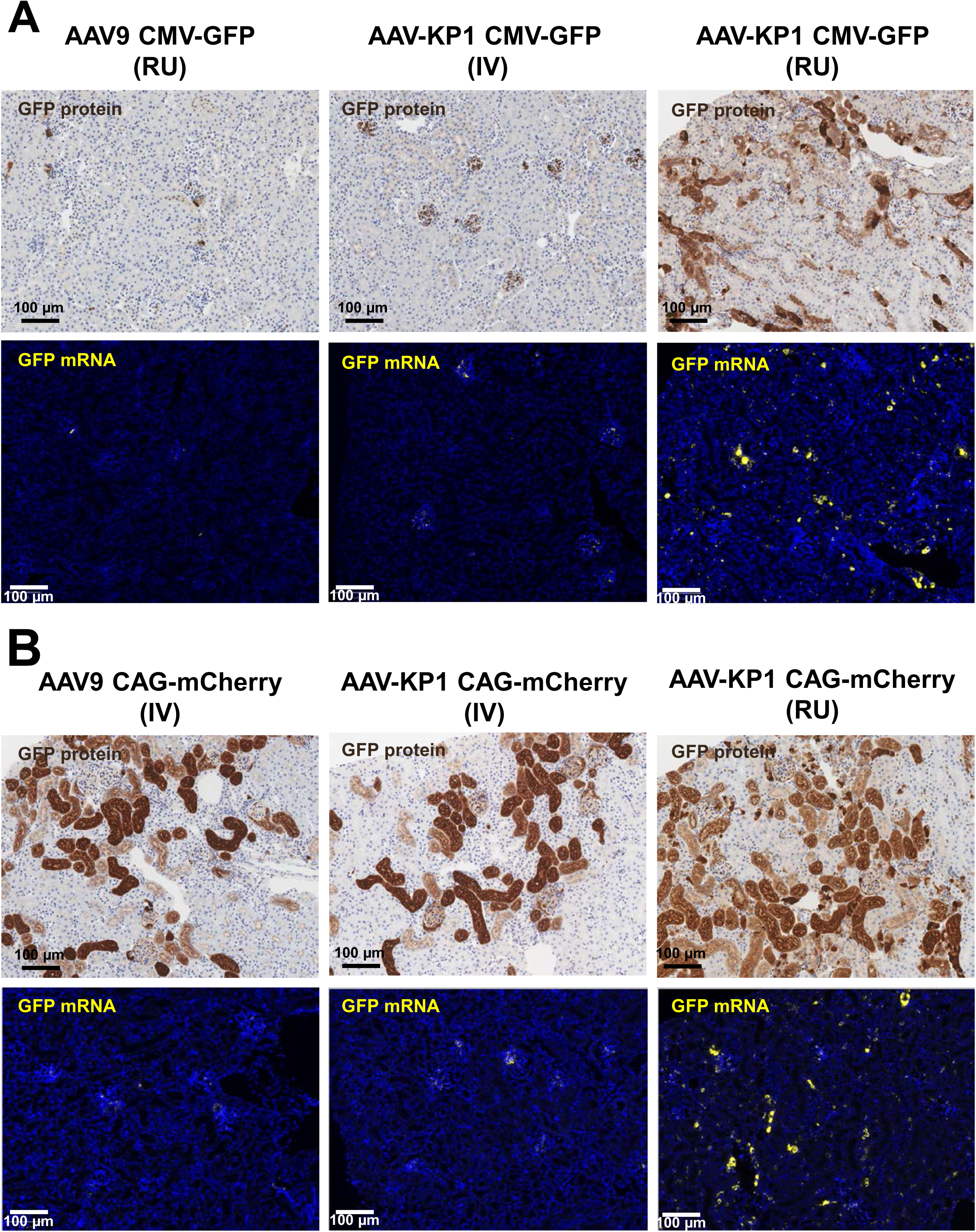
Comparison of transgene detection at protein and transcript level in the injected kidney. Six-week-old mice received 1 × 10¹⁰ vg/g of AAV-KP1 or AAV9 vectors carrying either GFP or mCherry under the CMV or the CAG promoter, delivered into the left renal pelvis (RU) or intravenously (IV). Mice were euthanised two weeks later. All panels show sections of the injected left kidney. **(A)** GFP protein detected by immunohistochemistry. **(B)** GFP transcripts detected by RNA fluorescence in situ hybridisation (FISH). **(C)** mCherry protein detected by immunohistochemistry. **(D)** mCherry transcripts detected by RNA FISH. Boxed regions are shown at higher magnification. Scale bars are indicated in each image.

Taken together, these results indicate that most of the transgenic protein detected in the non-injected kidney of locally administered animals, and in the kidneys of intravenously administered animals, does not arise from AAV-mediated transgene expression in those cells. Furthermore, the protein distribution pattern resembles that expected from kidney reabsorbed proteins like albumin or ApoA1. We therefore assessed mCherry presence in the serum of mice administered AAV9 CAG-mCherry intravenously, AAV-KP1 CAG-mCherry retro-ureterally, or PBS. mCherry was detected by ELISA in the serum of all mice receiving AAVs, at lower levels in the animals dosed retro-ureterally (Figure 5D). Notably, within this group, the three mice with the highest serum mCherry levels were also the three mice with the highest hepatic mCherry mRNA and renal mCherry protein levels (Figure 5A, Supplementary figure 8). This suggest that the recombinant proteins expressed in the liver reaches the circulation, by a mechanism that remains to be elucidated since they are intracellular proteins, and are subsequently taken up by the kidney.

## DISCUSSION

This study identifies AAV-KP1, AAV-KP2 and AAV-KP3 as AAV serotypes with strong and consistent tropism for the renal epithelium following retro-ureteral administration, and shows that this tropism is largely retained across promoters (KSPC and CMV). It also shows that a substantial fraction of the transgene protein detected in the proximal tubule after both local and systemic delivery reflects reabsorption of circulating transgene product rather than local transduction. Together, these findings have direct implications for how renal AAV transduction should be measured and interpreted, both in this work and in the broader field of kidney-directed gene therapy.

The barcoded library screen cleanly separated renal-tropic from hepatotropic serotypes selecting four capsids enriched in the retro-ureterally injected kidney (AAV-KP1, AAV-KP2, AAV-KP3, AAV.KK04) that were not among the serotypes enriched in the liver, and vice versa. This inverse relationship suggests that the structural determinants governing renal epithelial entry after direct pelvic delivery are largely non-overlapping with those governing hepatic uptake from the circulation, which is consistent with the fact that our administration route bypasses the glomerular filtration barrier and vascular hepatic uptake altogether. Nonetheless, the liver is still highly transduced when these capsids are administered retro-ureterally. Interestingly, intravenous administration of AAV-KP1 favoured hepatic transduction, indicating that a fraction of the vector escapes the injected kidney and reaches the systemic circulation. Conversely, intravenous administration of AAV-KP1 favoured hepatic over renal transduction, indicating that the route of administration is itself a determinant of apparent AAV tropism. Several studies have shown that the administration route affects AAV tropism, probably because the proteins that interact with the capsid, and thereby modulate its transduction capacity, differ with the biological compartment in which the vector is placed (33,34). However, we cannot exclude that the reduced hepatic transduction observed when AAV-KP1 is administered retro-ureterally simply reflects a smaller effective dose reaching the liver, as a consequence of vector particles being retained in the administered kidney.

The exclusion of AAV.KK04 from further characterization, based on its weaker renal transduction and reduced transgene expression relative to the other three candidates, reflects a prioritization rather than a lack of renal tropism, and this serotype may merit revisiting in contexts where its distinct profile is advantageous.

Among AAV-KP1, AAV-KP2 and AAV-KP3, transgene expression was similar despite differing genome copy numbers most notably the higher hepatic genome copy numbers but lower genome-normalized hepatic expression of AAV-KP2. This indicates that raw biodistribution data alone do not reliably predict functional transduction, and that normalization to genome copy number is necessary for a meaningful comparison of serotype potency. The sex-dependent expression pattern we observed with the AAV-KPs, with higher hepatic expression in females and higher renal (left kidney) expression in males, is a notable and, to our knowledge, underappreciated variable in renal AAV gene therapy studies. Sex-dependent differences in hepatic AAV transduction, as observed here in AAV9-treated mice, are well documented and are typically linked to hormonal control of receptor expression or intracellular processing (35,36). Our findings extend these sex-specific variations to the kidney, underscoring that biological sex should be incorporated as a deliberate design variable rather than a confounding factor in future dosing and efficacy studies. Because this analysis pooled all AAV-injected groups and was not powered a priori for a sex comparison, the magnitude of the effect should be confirmed in dedicated experiments.

Analysis of cellular tropism in the treated kidney revealed robust transduction of the collecting duct, distal convoluted tubule, proximal tubule, and Bowman’s capsule. These results reveal a broader pattern of renal epithelial transduction than that reported by other authors, who observed predominantly proximal tubular transduction with different AAV serotypes, including AAV-KP1 (20–22). Our study differs from those reports in several respects, including the serotypes examined and, in some cases, the use of systemic rather than local administration; as discussed below, the readouts used to score transduction also differ. AAV-KP1 and AAV-KP3 targeted a broad array of nephron segments, which is of direct therapeutic interest for monogenic renal disorders with segment-specific pathology, such as polycystic kidney disease. Beyond confirming renal tropism, the comparison of the KSPC and CMV promoters revealed wider renal cell-type coverage with CMV, extending transduction to interstitial and podocyte populations, whereas Bowman’s capsule cells were detected only with KSPC. The absence of GFP signal in the thick ascending limb with both promoters leaves open whether this reflects a true transduction boundary or a gap in promoter activity. These observations highlight the need for panels of renal promoters when evaluating renal AAV capsids, since cell-type coverage inferred from a single promoter will systematically underestimate the true transduction profile. The unexpectedly high hepatic activity of the KSPC promoter further underscores the need for genuinely renal-specific promoters in the AAV genome context. In fact, the specificity of promoter fragments characterized in other settings, such as transgenic mice or plasmid transfection, cannot be assumed to be retained in an AAV vector.

The most significant finding of this study, however, is the evidence that transgene protein detected in the renal proximal tubule after both local and systemic administration substantially reflects reabsorption of circulating transgene product rather than local transduction, and that this is not merely an assay artifact confined to one construct or serotype. Four independent lines of evidence converge on this conclusion. First, abundant GFP or mCherry protein coexisted with low or absent transgene mRNA in renal samples from AAV9-treated and intravenously administered AAV-KP1-treated mice, and in the contralateral kidney when AAV-KP1 was administered into the left kidney; such a discordance is difficult to reconcile with genuine transduction, since a transduced cell should contain both transcript and protein. Second, the apical distribution of the fluorescent protein in proximal tubule cells mirrors the well-established pattern of megalin/cubilin-mediated reabsorption of filtered proteins such as albumin and apolipoprotein A-I (37–40). Third, mCherry was detectable in the serum of AAV-treated animals and, within the locally treated AAV-KP1 group, the animals with the highest serum mCherry were also those with the highest hepatic mCherry mRNA and the highest renal mCherry protein. This pattern is consistent with hepatic secretion into the circulation, glomerular filtration and downstream tubular reabsorption, rather than with direct transduction of the distal organ. Fourth, RNA-FISH confirmed that transgene mRNA was essentially absent from the epithelial regions showing strong protein signal in the non-transduced or systemically exposed kidneys, whereas mRNA, genome copy number and protein signal were concordant in the left kidney after retro-ureteral administration of AAV-KP1.

Two aspects of this phenomenon deserve comment. The first is how intracellular reporters reach the circulation at all. Neither GFP nor mCherry carries a secretory signal peptide, so their presence in serum most likely reflects passive release from transduced cells through physiological cell turnover, vector-or transgene-related cytotoxicity, or membrane damage associated with the procedure; given the sensitivity of the ELISA and IHC readouts, low-level leakage from the large mass of transduced hepatocytes would be sufficient. Although the mechanism remains to be defined, its practical consequence is clear: any transgene product that enters the circulation and crosses the glomerular filtration barrier can be reabsorbed and misread as local transduction.

The second aspect is that the magnitude of this interference is transgene-dependent. Under physiological conditions, protein size limits passage across the glomerular filtration barrier, with free filtration occurring for proteins up to approximately 60-70 kDa (41); albumin (∼66-69 kDa) is typically regarded as the practical cut-off, being largely retained under normal conditions. Both GFP and mCherry (∼27 kDa) are therefore well below this limit and freely filtered, yet they affected the assessment of renal transduction to different extents and robustness, indicating that additional properties of the transgene product, such as net charge, conformational stability, and affinity for the megalin/cubilin endocytic receptors, shape how much filtered protein is retained in the tubular epithelium. The susceptibility of a given reporter or therapeutic protein to this artifact therefore cannot be assumed and should be established empirically, with secreted therapeutic proteins below the filtration cut-off expected to be particularly susceptible.

Previous studies in mice have characterized AAV serotypes using systemic administration, or renal administration via the renal vein or the retro-ureteral route, with some observations similar to ours but also several differences that may be attributable to transgene reabsorption. Earlier studies comparing the renal tropism of AAV9 and AAV-KP1 reported that only locally administered AAV-KP1 delivered vector genomes efficiently to the treated kidney (21). In those studies, however, transgene expression was assessed solely by mCherry visualisation in renal sections, which showed a predominantly proximal tubular signal. Using GFP as the reporter, we instead detect AAV-KP1 transduction across additional segments of the renal epithelium. A separate study delivered two vectors, including AAV9, intravenously (22). Renal transduction was again assessed primarily by mCherry visualisation, showing a predominantly proximal tubular pattern, but neither transgene mRNA nor vector genome copies were systematically quantified in the organs examined. Where copy numbers were measured, transgene protein differed across serotypes at comparable vector genome loads. Intraperitoneal administration of AAV1 has also been proposed as a route to renal transduction, showing GFP in the renal epithelium and in renal cysts, along with detectable transgene mRNA and AAV genomes in the kidney (42). Benchmarking these studies against local renal administration routes such as ours would require renal transduction to be reported relative to hepatic transduction, since it is the renal-to-hepatic ratio, rather than the renal signal alone, that determines both the specificity of a delivery strategy and the likelihood that a renal protein signal is of extra-renal origin.

This reabsorption phenomenon has practical consequences for how renal AAV biodistribution and efficacy studies are designed and interpreted. Protein-level readouts alone are insufficient to establish local transduction in the kidney, and particularly in the proximal tubule, because reabsorbed protein can closely mimic the appearance of true transduction; this holds even for a protein retained intracellularly rather than actively secreted, provided that AAV-transduced cells elsewhere in the body release it into the circulation. Our data therefore suggest that transgene mRNA detection (by RNA-FISH and RT-qPCR) together with vector genome quantification should be considered a necessary complement to protein readouts, rather than an optional confirmatory step, whenever transduction of the kidney is claimed. RNA-FISH is very informative because it preserves spatial information and therefore allows the cell type producing the transcript to be identified, which is precisely the information lost when protein staining is interpreted on its own. In studies using unilateral local delivery, the contralateral kidney provides a convenient internal control for reabsorption, since it is exposed to the same circulating protein but receives minimal vector.

Applying these criteria to our own data allows two distinct phenomena to be separated. The consistent detection of both protein and mRNA in glomeruli, for GFP and for mCherry, across essentially all treated groups, including systemically injected mice, indicates that glomerular cells, most likely podocytes, although mesangial cells cannot be excluded (21), and consistent with the AAV9 tropism data, are genuinely and efficiently transduced through the vasculature. In contrast, the retro-ureterally administered kidney showed genuinely broad epithelial transduction with AAV-KP1, AAV-KP2 and AAV-KP3, confirmed by concordant mRNA and protein and by genome copy numbers approaching hepatic levels, whereas the contralateral kidney and the kidneys of systemically treated animals showed a glomerulus-dominated mRNA pattern with superimposed tubular protein reabsorption. This distinction is central to correctly attributing renal epithelial transduction to the AAV-KP capsids specifically and to the retro-ureteral route specifically, rather than to a generic property of these AAV vectors. It also clarifies why AAV9, despite genuine access to the kidney via retro-ureteral delivery, achieved substantially lower renal genome copy numbers and transduction than AAV-KP1 or AAV-KP3 under the same conditions, reinforcing that renal epithelial entry after direct pelvic exposure is capsid-dependent and not simply a function of local vector concentration.

While this work was in preparation, Sairavi and colleagues independently reported a closely related phenomenon in mice and in marmoset (43). Their conclusions agree with ours, in that monomeric fluorescent reporters released into the circulation from transduced organs are filtered and reabsorbed by the proximal tubule, where the resulting protein signal can be mistaken for local transduction. This independent agreement strengthens both studies. Their work shows that megalin is directly responsible, by conditional deletion of the receptor in the proximal tubule, and that the phenomenon also seems to occurs in non-human primates. Our data add two complementary elements. First, we detect transgene transcript at cellular resolution by RNA-FISH, so the cell that produces the transcript is identified directly rather than inferred from the subcellular distribution of the protein. Second, unilateral retro-ureteral delivery provides a control within the same animal, since the contralateral kidney is exposed to the same circulating protein but receives almost no vector, so reabsorption and true transduction can be distinguished in one animal with the same construct.

Taken together, this work (1) identifies AAV-KP1, AAV-KP2 and AAV-KP3 as strong candidate vectors for renal epithelial gene delivery via retro-ureteral administration, with transduction efficiency in the target kidney approaching that of the liver; (2) highlights promoter choice, biological sex and multi-promoter characterization as variables that materially affect the interpretation of renal AAV biodistribution data; and (3) establishes that reabsorption of circulating transgene product by the proximal tubule can closely mimic local transduction at the protein level, so that confirmation of transgene mRNA presence is required before renal transduction is claimed. Further work extending these observations to disease models, and assessing the durability and functional consequences of AAV-KP1-and AAV-KP3-mediated transduction of the diverse epithelial segments identified here, will be needed to translate this renal-tropic platform toward therapeutic applications in monogenic and acquired kidney diseases.

## ABBREVIATIONS

AAV: Adeno-associated virus
cDNA: Copy DNA
CKD: Chronic kidney disease
CMV: Cytomegalovirus
gDNA: Genomic DNA
GFP: Green Fluorescent Protein
mRNA: Messenger RNA
IF: Immunofluorescence
IHC: Immunohistochemistry
IV: Intravenous
kDa: Kilodaltons
KSPC: Kidney-specific cadherin
LK: Left Kidney
PCR: Polymerase Chain Reaction
PEI: Polyethylenimine
RK: Right kidney
RNA-FISH: RNA fluorescence in situ hybridisation
RT-qPCR: reverse transcription quantitative polymerase chain reaction
RU: Retro-ureteral

## Data Availability Statement

Sequencing data generated in this study was deposited at NCBI SRA under BioProject ID PRJNA1552391. The datasets supporting the conclusions of this study are included within the article and its supplementary data file.

## ACKNOWLEDGMENTS

We gratefully acknowledge the support of the CIMA Genomics Platform, particularly Dr. P. Marchese, and the CIMA Morphology Platform, especially L. Guembe, for their valuable technical assistance. We also sincerely thank the Centro de Investigación Médica Aplicada (CIMA) and all colleagues from the RNA and DNA Medicine Division for their support and collaboration. Special thanks go to the members of the Rare Disease Gene Therapy Program, particularly D. Herrero and Dr A. Llanos-Ardaiz, for their assistance with various experimental procedures.

## AUTHOR CONTRIBUTIONS

Conceptualization, R.A., G.G-A.; data curation, M.B-A, S.M.; formal analysis, M.B-A., S.M.; funding acquisition, G.G-A., R.A.; investigation, S.M., N.S-G.; methodology, S.M., N.S-G., L.N., B.C., A.V., M.M.; project administration, R.A., G.G-A.; resources, R.A., L.L.; software, M.B-A.; supervision, N.Z., G.G-A., R.A; validation, M.B-A., S.M., N.S-G.; visualization, M.B-A., S.M, N.S-G. writing – original draft, R.A.; writing – review & editing, all authors.

## FUNDING

Rafael Aldabe disclose support for the research of this work from Spain Ministry of Science Innovation and Universities (PID2022-143327OB-I00, PID2025-176597OB-I00). Rafael Aldabe and Gloria Gonzalez-Aseguinolaza report financial support provided by Navarra Government General Directorate of Industry Energy and Strategic Projects S3 (0011-1411-2024-000121). Sergio Milagros disclose support from by Navarra Government General Directorate of Industry Energy and Strategic Projects S3 (0011-1408-2020-000028).

## ETHICAL APPROVAL

All animal experiments were in accordance with Directive 2010/63/EU and Spanish Royal Decree 53/2013 (as amended by RD 1386/2018 and RD 118/2021), with approval from the Animal Experimentation Ethics Committee of the University of Navarra and authorization from the Government of Navarra (069-20 and 117-25).

## COMPETING INTERESTS

G.G-A. is co-founder and shareholder of Vivet Therapeutics and Amets Biotechnology. L.L. is co-founder, Chief Technology Officer and shareholder of of Nionyx Bio.

---

**Figure 5. Renal and hepatic transduction after administration of AAV-KP1 or AAV9 CAG-mCherry vectors.**

Six-week-old mice received 1 × 10¹⁰ vg/g of the AAV-KP1 vector carrying mCherry under the CAG promoter, delivered either into the left renal pelvis (RU) or intravenously (IV), or of the AAV9 CAG-mCherry vector delivered intravenously. Mice were euthanised two weeks later to quantify vector genomes and mCherry transgene expression. **(A)** mCherry-positive cells detected by immunohistochemistry in sections of the left (upper sections) and right (lower sections) kidneys. Boxed regions are shown at higher magnification. Scale bars are indicated in each image. **(B)** Vector genome (vg) copies in liver and in both kidneys, measured by quantitative PCR. **(C)** mCherry transcript levels in liver and in both kidneys, measured by RT-qPCR. **(D)** Serum mCherry concentration, quantified by ELISA. Data are mean ± SD. Groups were compared using Tukey’s multiple comparisons test; numerical values indicate P values for each pairwise comparison.

