## Supplemental Figures for "RETRO-URETERAL DELIVERY OF AAV-KP VECTORS FOR RENAL GENE TRANSFER AND THE ASSESSMENT OF TRANSGENE REABSORPTION IN THE KIDNEY"

### Supplementary Figure 1

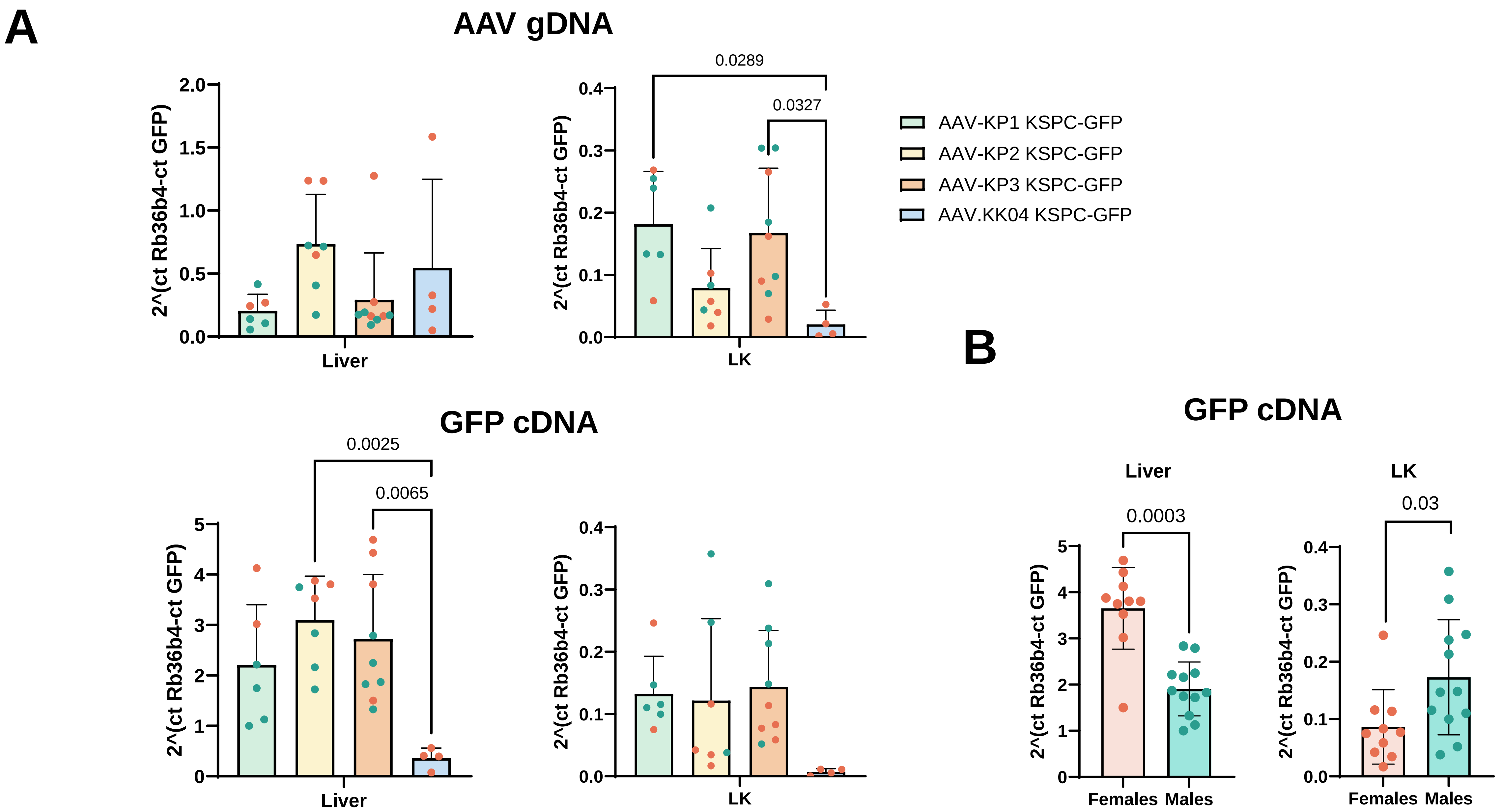

#### Supplementary Figure 2

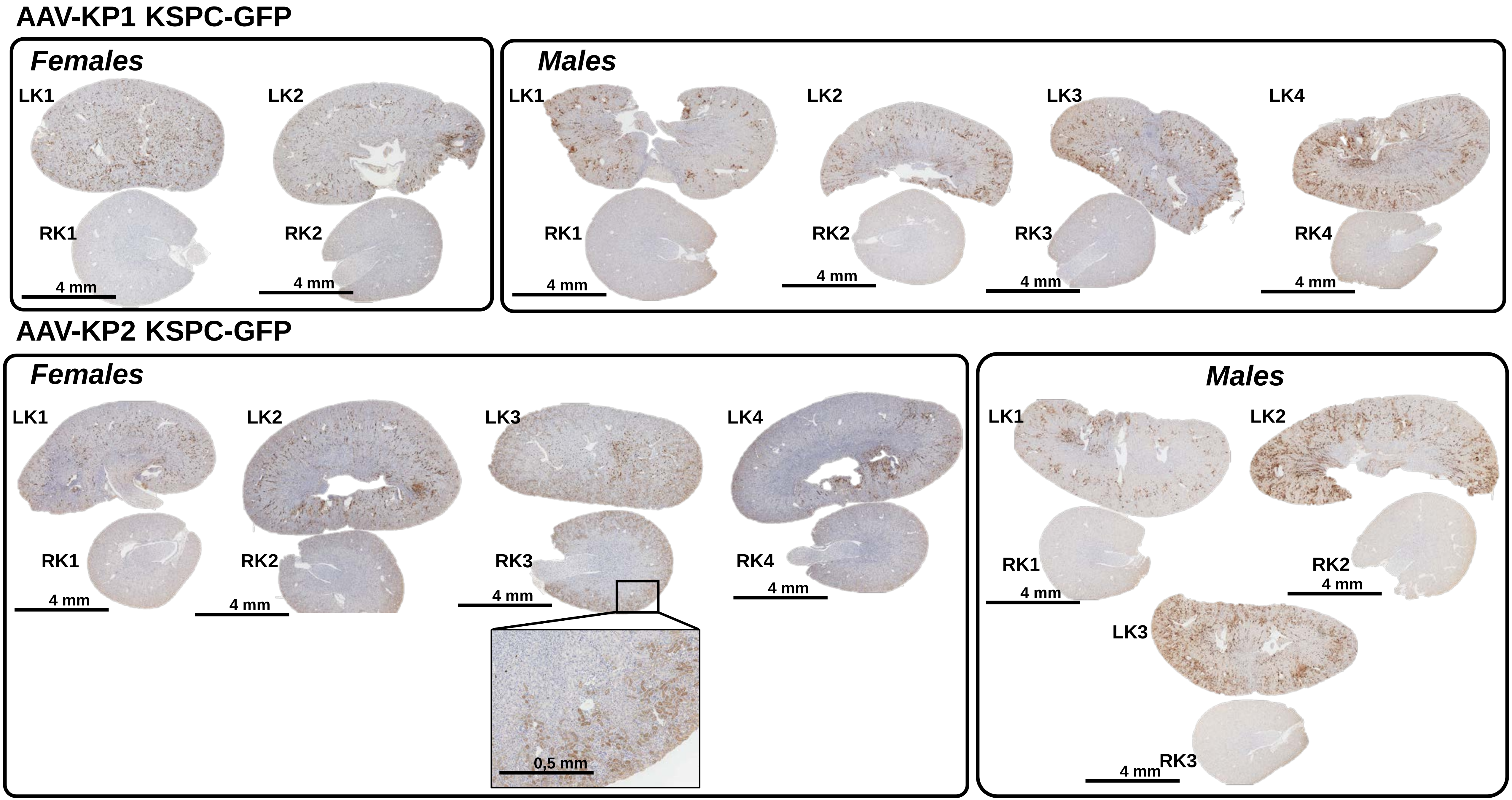

### Supplementary Figure 2

#### AAV-KP3 KSPC-GFP

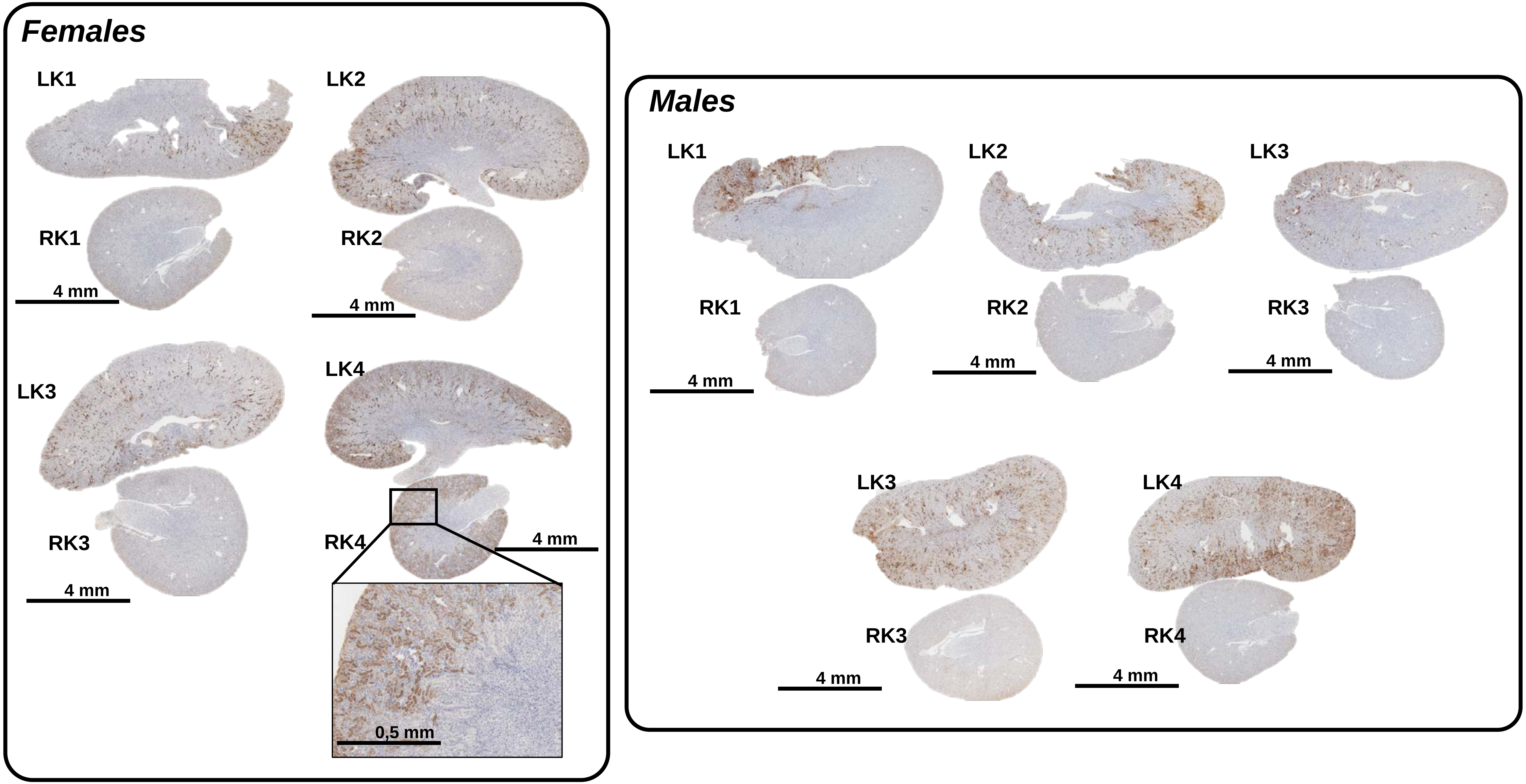

#### Supplementary Figure 3

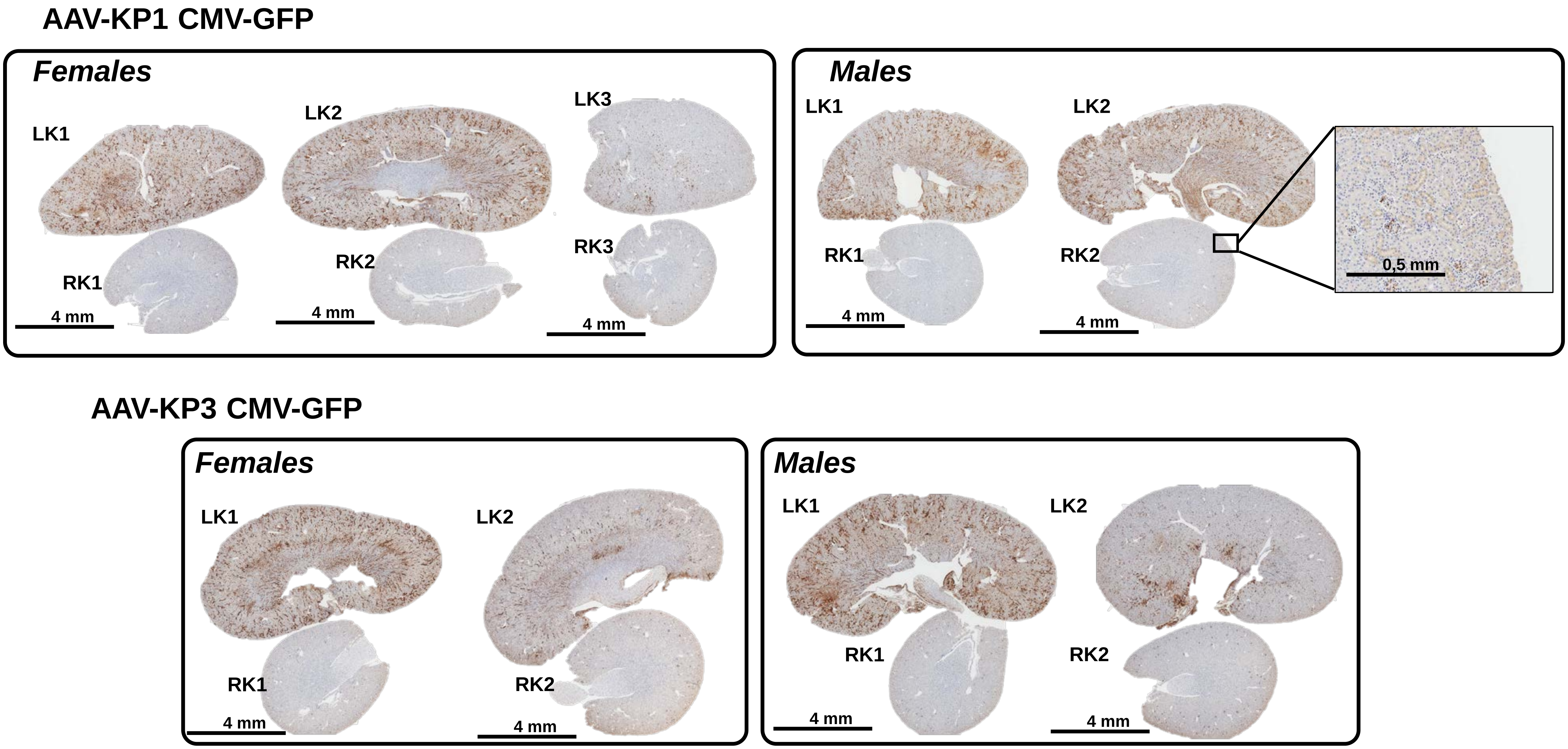

### Supplementary Figure 3

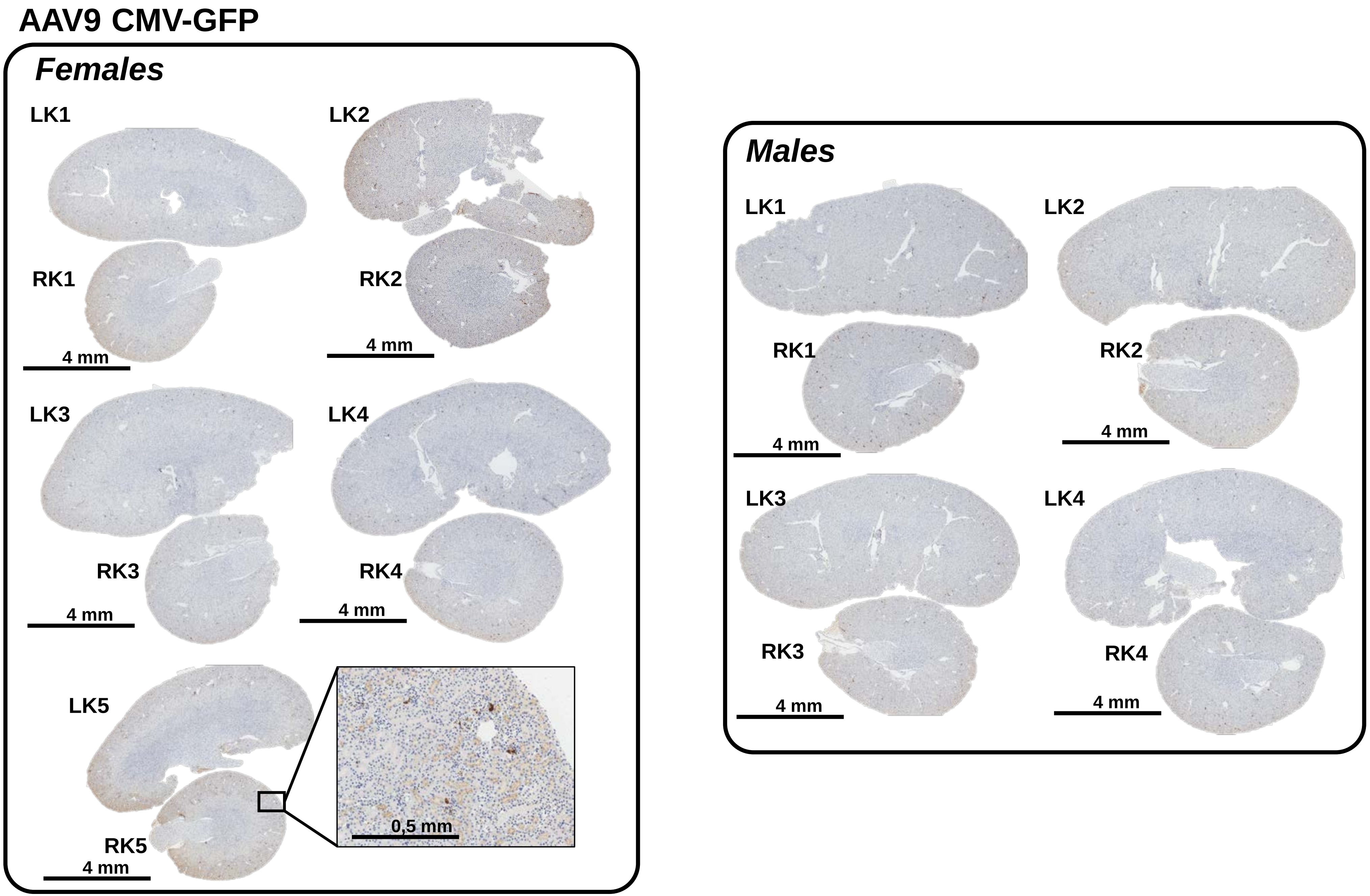

### Supplementary Figure 4

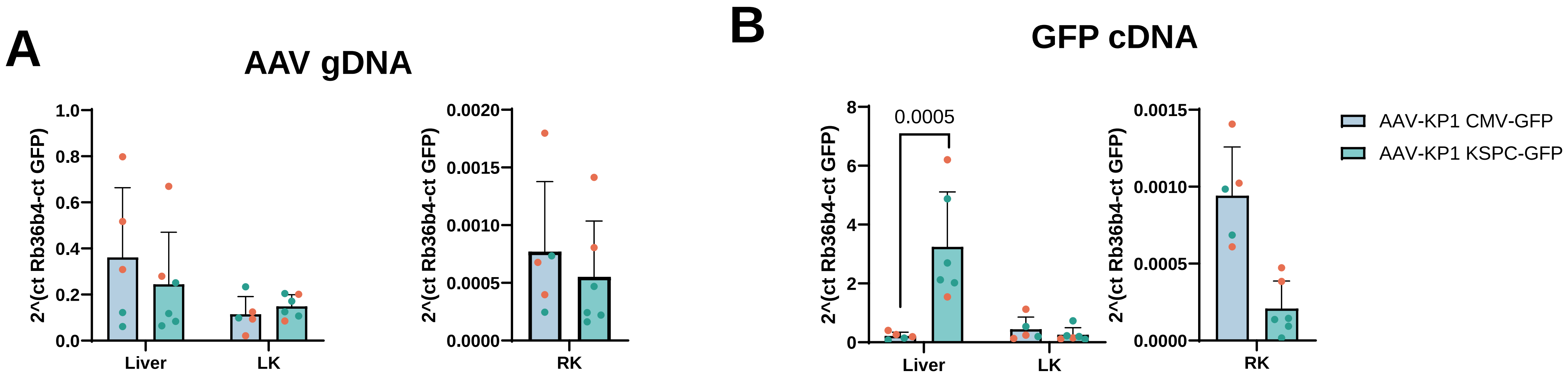

### Supplementary Figure 5

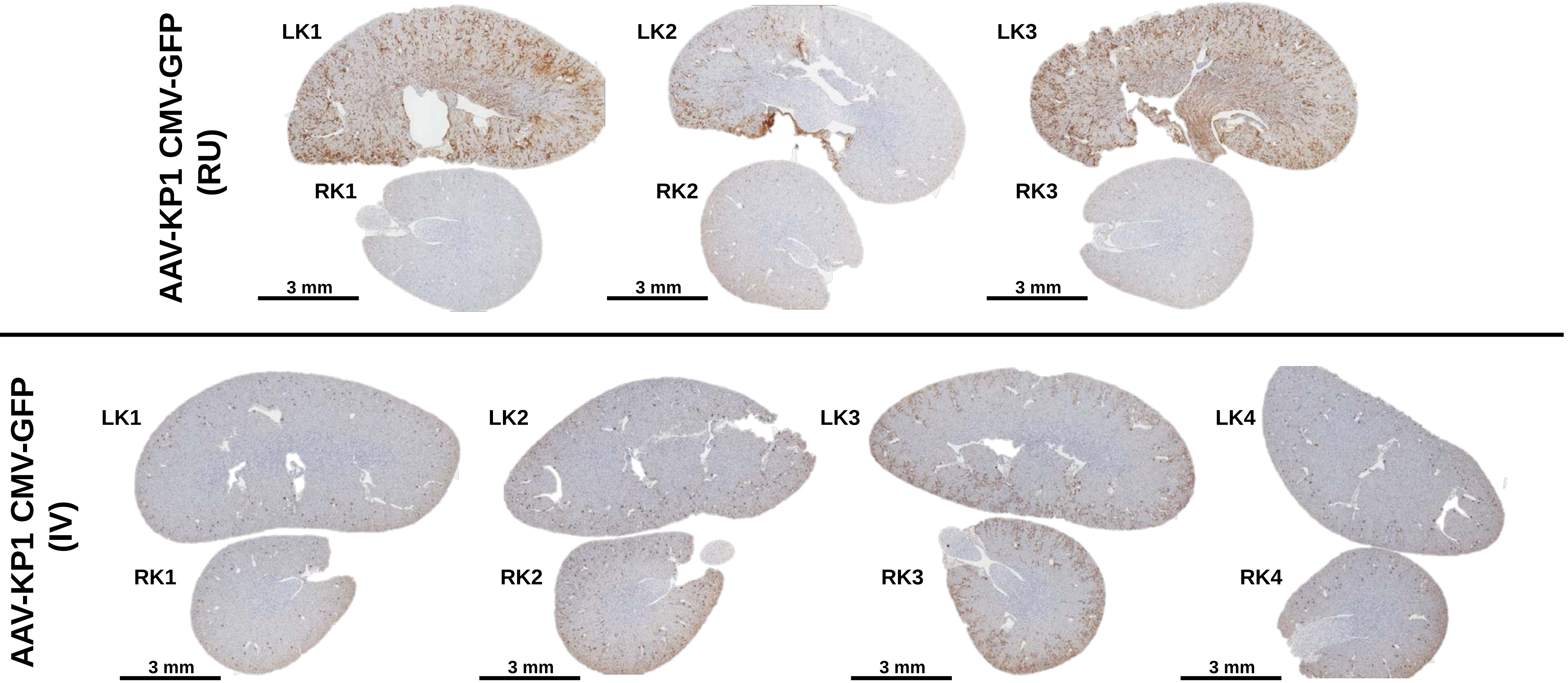

### Supplementary Figure 6

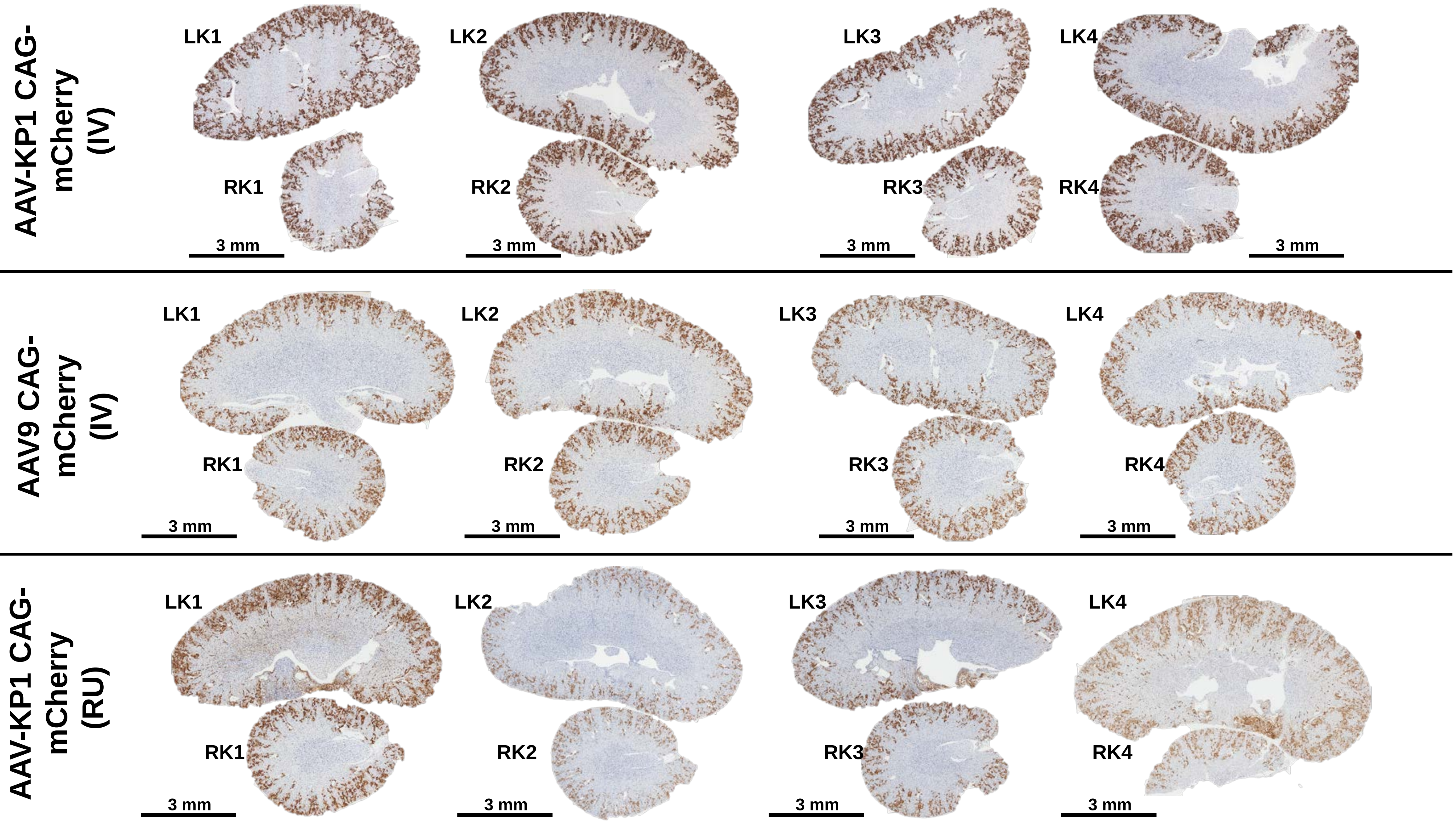

### Supplementary Figure 7

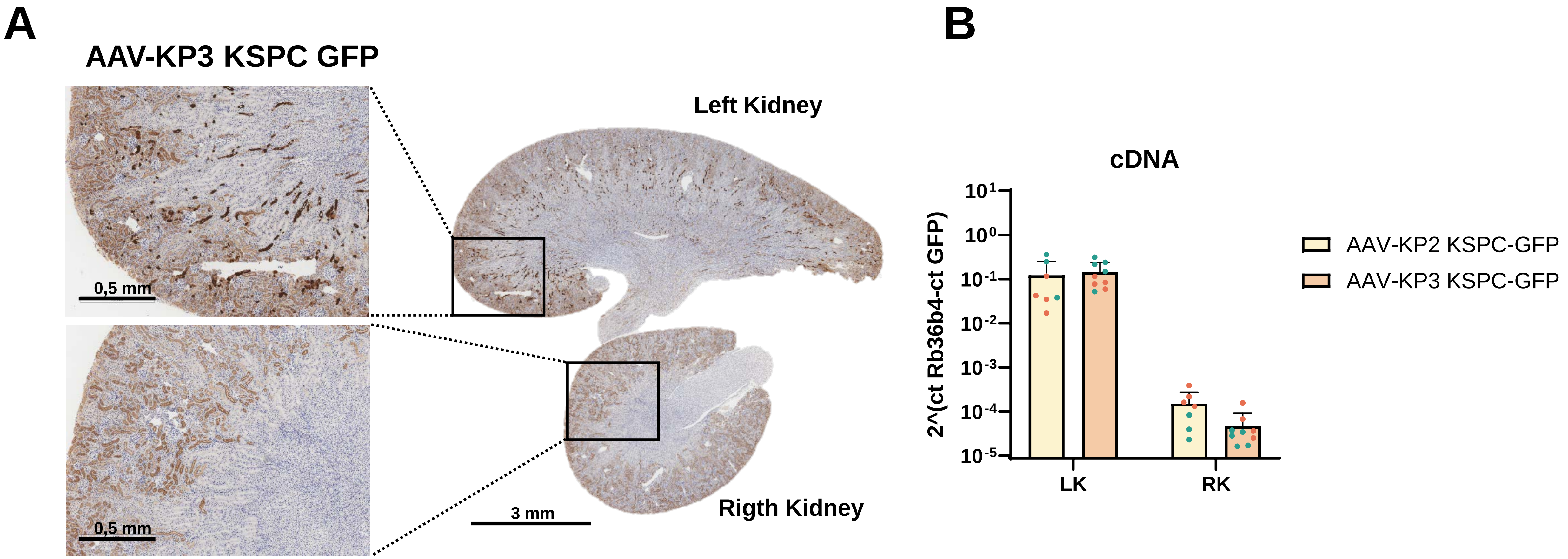

### Supplementary Figure 8

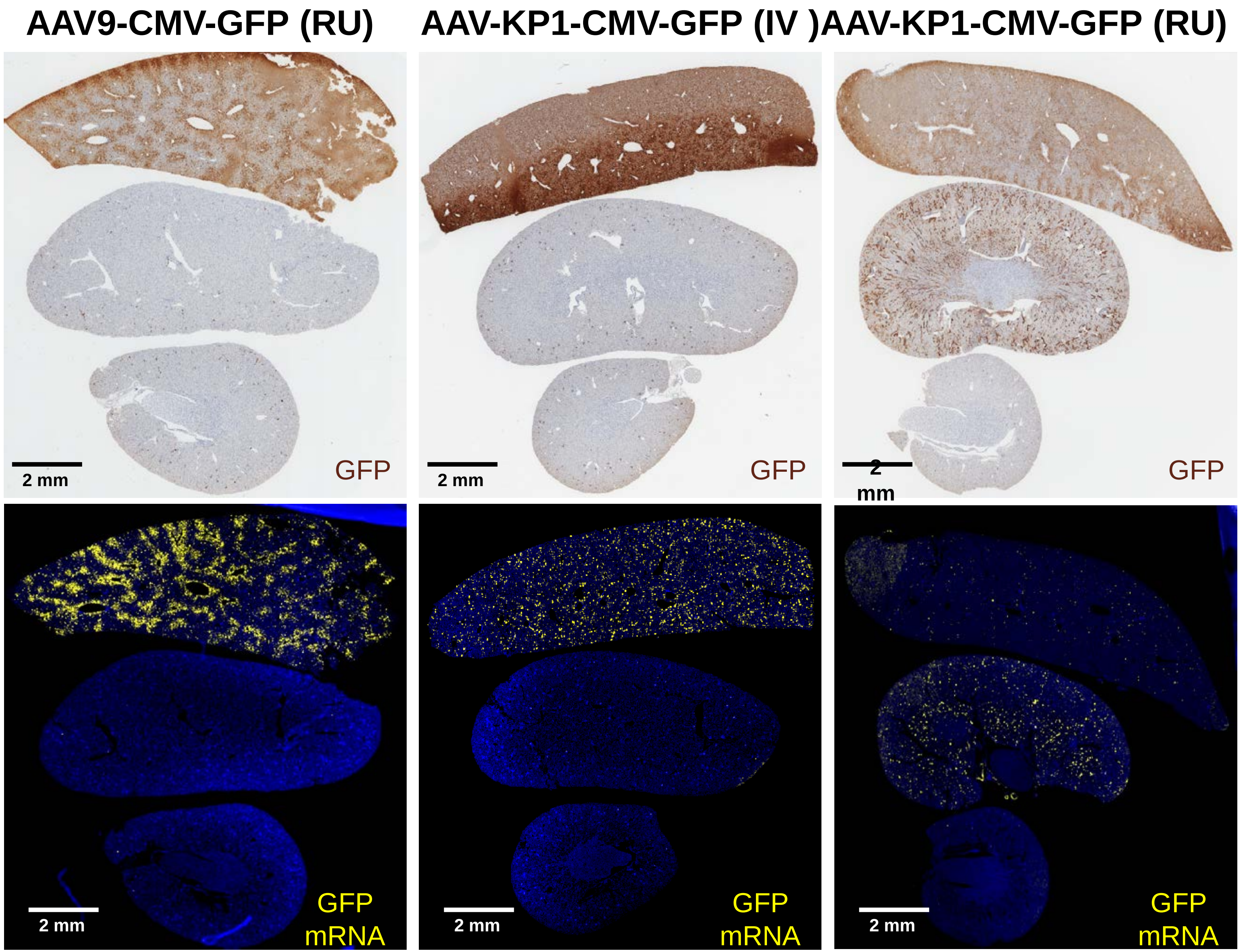

### Supplementary Figure 9

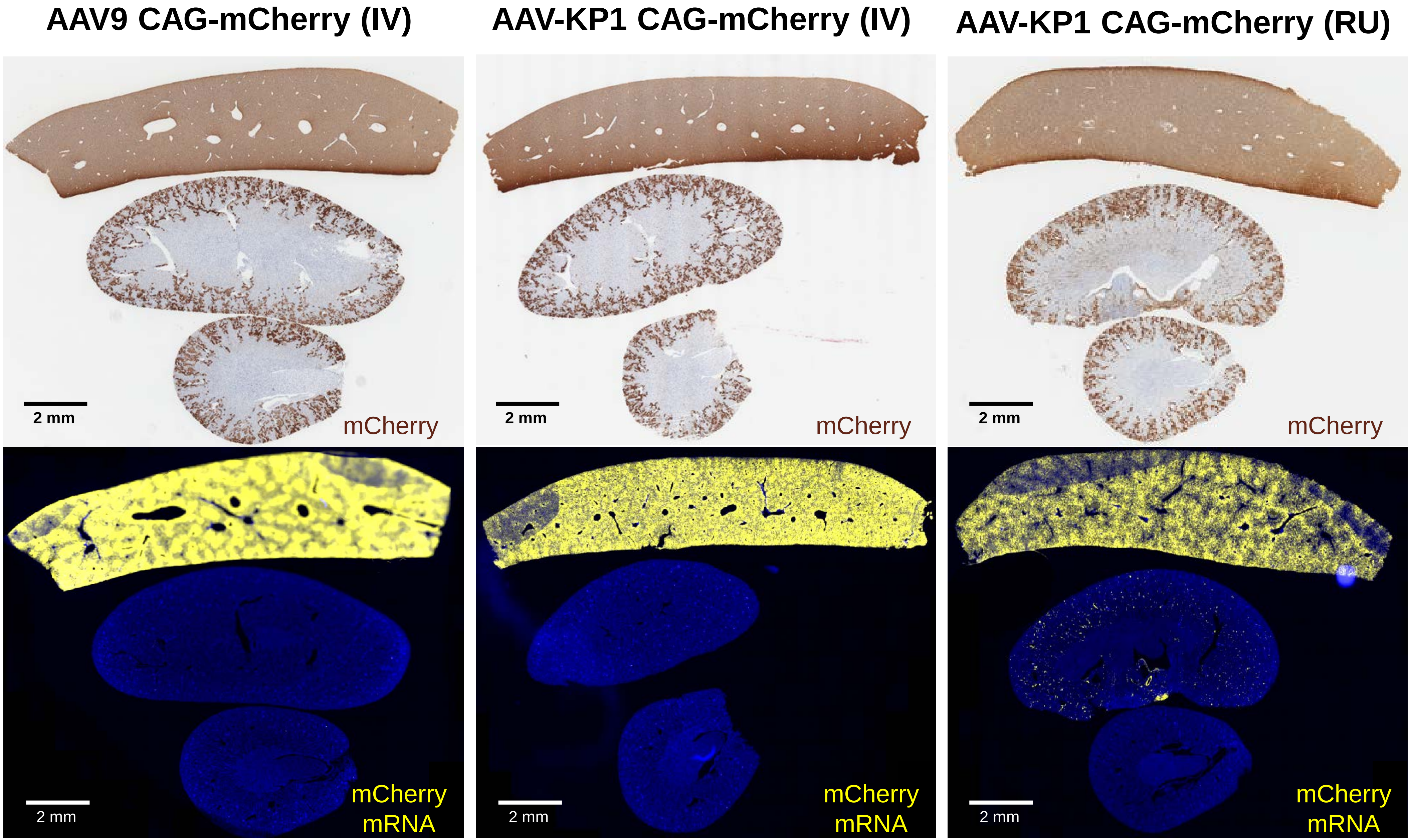

#### ADDITIONAL INFORMATION

#### SUPPLEMENTARY FIGURES LEGENDS

**Supplementary Figure 1. Renal and hepatic transduction efficiency of AAV-KP1, AAV-KP2, AAV-KP3 and AAV.KK04 after retro-ureteral administration.** Six-week-old mice received 1 × $10^{10}$ vg/g of AAV-KP1, AAV-KP2, AAV-KP3 or AAV.KK04 vectors carrying GFP under the KSPC promoter, injected into the left renal pelvis. Mice were euthanised two weeks later to quantify vector genomes and GFP transgene expression. **(A)** Vector genome (vg) copies and GFP transcript levels in the liver and left kidney two weeks after administration, measured by quantitative PCR and RT-qPCR. **(B)** GFP transcript levels in the liver and left kidney two weeks after administration of AAV-KP1, AAV-KP2 or AAV-KP3, quantified by RT-qPCR. Animals were pooled by sex (males, green symbols; females, orange symbols).

**Supplementary Figure 2. Renal transduction by AAV-KP1, AAV-KP2 and AAV-KP3 following retro-ureteral administration.** Six-week-old mice received 1 × $10^{10}$ vg/g of AAV-KP1, AAV-KP2 or AAV-KP3 encoding GFP under the KSPC promoter, injected into the left renal pelvis. Two weeks later, mice were euthanised and GFP expression was quantified. Immunohistochemical detection of GFP-positive cells in sections of the left (LK) and right (RK) kidneys of the males and females animals that showed detectable transduction of the left kidney. Boxed regions are shown at higher magnification; scale bars as indicated.

**Supplementary Figure 3. Renal transduction by AAV-KP1, AAV-KP2 and AAV9 following retro-ureteral administration.** Six-week-old mice received 1 × $10^{10}$ vg/g of AAV-KP1, AAV-KP2 or AAV9 encoding GFP under the CMV promoter, injected into the left renal pelvis. Two weeks later, mice were euthanised and GFP expression was quantified. Immunohistochemical detection of GFP-positive cells in sections of the left (LK) kidneys of the males and females animals that showed detectable transduction of the left kidney. Boxed regions are shown at higher magnification; scale bars as indicated.

**Supplementary Figure 4. Renal and hepatic transduction efficiency of AAV-KP1 after retro-ureteral administration.** Six-week-old mice received 1 × $10^{10}$ vg/g of AAV-KP1 vector carrying GFP under the KSPC or the CMV promoter, injected into the left renal pelvis. Mice were euthanised two weeks later to quantify vector genomes and GFP transgene expression. **(A)** Vector genome (vg) copies and **(B)** GFP transcript levels in the liver and left kidney two weeks after administration, measured by quantitative PCR and RT-qPCR.

**Supplementary Figure 5. Renal transduction by AAV-KP1 following retro-ureteral and systemic administration.** Six-week-old mice received 1 × $10^{10}$ vg/g of AAV-KP1, encoding GFP under the CMV promoter, injected into the left renal pelvis. Two weeks later, mice were euthanised and GFP expression was quantified. Immunohistochemical detection of GFP-positive cells in sections of the left and right kidneys of the males that showed detectable transduction of the left kidney. Scale bars as indicated.

**Supplementary Figure 6. Renal transduction efficiency of the AAV-KP1 CAG-mCherry vector after systemic or retro-ureteral administration and AAV9 CAG-mCherry vector after systemic administration.** Six-week-old mice received 1 × $10^{10}$ vg/g of the AAV-KP1 or AAV9 vector carrying mCherry under the CAG promoter, delivered either into the left renal pelvis (RU) or intravenously (IV). Mice were euthanised two weeks later to quantify mCherry transgene expression. mCherry-positive cells detected by immunohistochemistry in sections of the right (lower kidney) and left (upper kidney) kidneys from all treated animals.

**Supplementary Figure 7. Comparison of GFP protein detection and mRNA expression after retro-ureteral administration of AAV-KP3 and AAV-KP2 vectors.** Six-week-old mice received 1 × $10^{10}$ vg/g of the AAV-KP3 or AAV-KP2 vector carrying GFP under the CMV promoter, delivered either into the left renal pelvis. Mice were euthanised two weeks later to quantify GFP transgene expression by IHC **(A)** and by RT-qPCR **(B)**.

**Supplementary Figure 8. Comparison of GFP detection at the protein and transcript level in the liver and both kidneys.** Six-week-old mice received 1 × $10^{10}$ vg/g of AAV-KP1 or AAV9 encoding GFP under the CMV promoter, delivered either into the left renal pelvis (RU) or intravenously (IV). Mice were euthanised two weeks later. **(A)** GFP protein detected by immunohistochemistry. **(B)** GFP transcripts detected by RNA fluorescence in situ hybridisation (FISH).

**Supplementary Figure 9. Comparison of mCherry detection at the protein and transcript level in the liver and both kidneys.** Six-week-old mice received 1 × $10^{10}$ vg/g of AAV-KP1 or AAV9 encoding mCherry under the CMV promoter, delivered either into the left renal pelvis (RU) or intravenously (IV). Mice were euthanised two weeks later. **(A)** mCherry protein detected by immunohistochemistry. **(B)** mCherry transcripts detected by RNA fluorescence in situ hybridisation (FISH).
